# Specialisation and plasticity in a primitively social insect

**DOI:** 10.1101/2020.03.31.007492

**Authors:** S. Patalano, A. Alsina, C. Gregorio-Rodriguez, M. Bachman, S. Dreier, I. Hernando-Herraez, P. Nana, S. Balasubramanian, S. Sumner, W. Reik, S. Rulands

## Abstract

Biological systems not only have the remarkable capacity to build and maintain complex spatio-temporal structures in noisy environments, they can also rapidly break up and rebuild such structures. How such systems can simultaneously achieve both robust specialisation and plasticity is poorly understood. Here we use primitive societies of *Polistes* wasps as a model system where we experimentally perturb the social structure by removing the queen and follow the re-establishment of the social steady state over time. We combine a unique experimental strategy correlating time-resolved measurements across vastly different scales with a theoretical approach. We show that *Polistes* integrates antagonistic processes on multiple scales to distinguish between extrinsic and intrinsic perturbations and thereby achieve both robust specialisation and rapid plasticity. The long-term stability of the social structure relies on dynamic DNA methylation which controls transcriptional noise. Such dynamics provide a general principle of how both specialization and plasticity can be achieved in biological systems.

**One Sentence Summary:** A primitive social insect simultaneously achieves specialisation and plasticity by integrating antagonistic dynamics on different scales.

**Highlights:**

- We employ a unique experimental approach correlating dynamics of societies, individuals, and epigenetic gene regulation
- A social insect simultaneously achieves specialisation and plasticity by integrating antagonistic processes on different spatial scales
- Regulation of population-level noise by DNA methylation ensures long-term stability of phenotypic specialisation

## Introduction

Biological systems have the remarkable capacity to build and maintain complex spatio-temporal structures. Such structures are often surprisingly robust in noisy environments and their formation relies on the integration of regulatory processes on vastly different spatial scales of organisation, from the molecular level to tissue or population-level feedback(*1*). While historically theoretical and experimental research has focussed on the processes underlying the formation of complex structures(*2–4*), in recent years it has become clear that biological systems also have the remarkable capacity to break up and rebuild these structures(*5*). As an example, colonies of social insects rely on the robust specialisation of individuals into distinct castes, such as queen and worker polyphenisms(*6*). Although such phenotypes can be stable over years in the face of environmental noise, individuals are capable of being phenotypically reprogrammed and perform different tasks from those which they performed initially (plasticity)(*7–9*). But how can biological systems achieve rapid plasticity to specific cues without specialised states being destabilised by noise?

To answer this question, we used a well-established model of phenotypic plasticity, the primitively social paper wasp *Polistes canadensis* (Fig. 1A) (*10–15*). Soon after the emergence of the foundress’ daughters (workers), a stable society of the paper wasp is established, with a single reproductive queen and 8-30 non-reproductive workers. If the queen dies (or is experimentally removed), the remaining workers can rapidly reprogram to generate a new queen, hence displaying strong phenotypic plasticity (*10, 16, 17*). We developed an experimental approach to follow the reprogramming dynamics simultaneously across different spatial scales: from population-level measurements, to physiological characterisation of the individuals and detailed molecular analyses of their brains using whole genome multi-omics (RNA-sequencing of the transcriptome and Bisulfite-sequencing of the methylome) (Fig. 1B). Specifically, we video-recorded all nests across the duration of the experiment and subsequently collected all individuals in each phase of the experiment for further dissection and sequencing (Fig. 1C). A unique feature of this approach is that it allows us to correlate dynamic measurements on different spatial scales at the individual level.

**Fig. 1.**
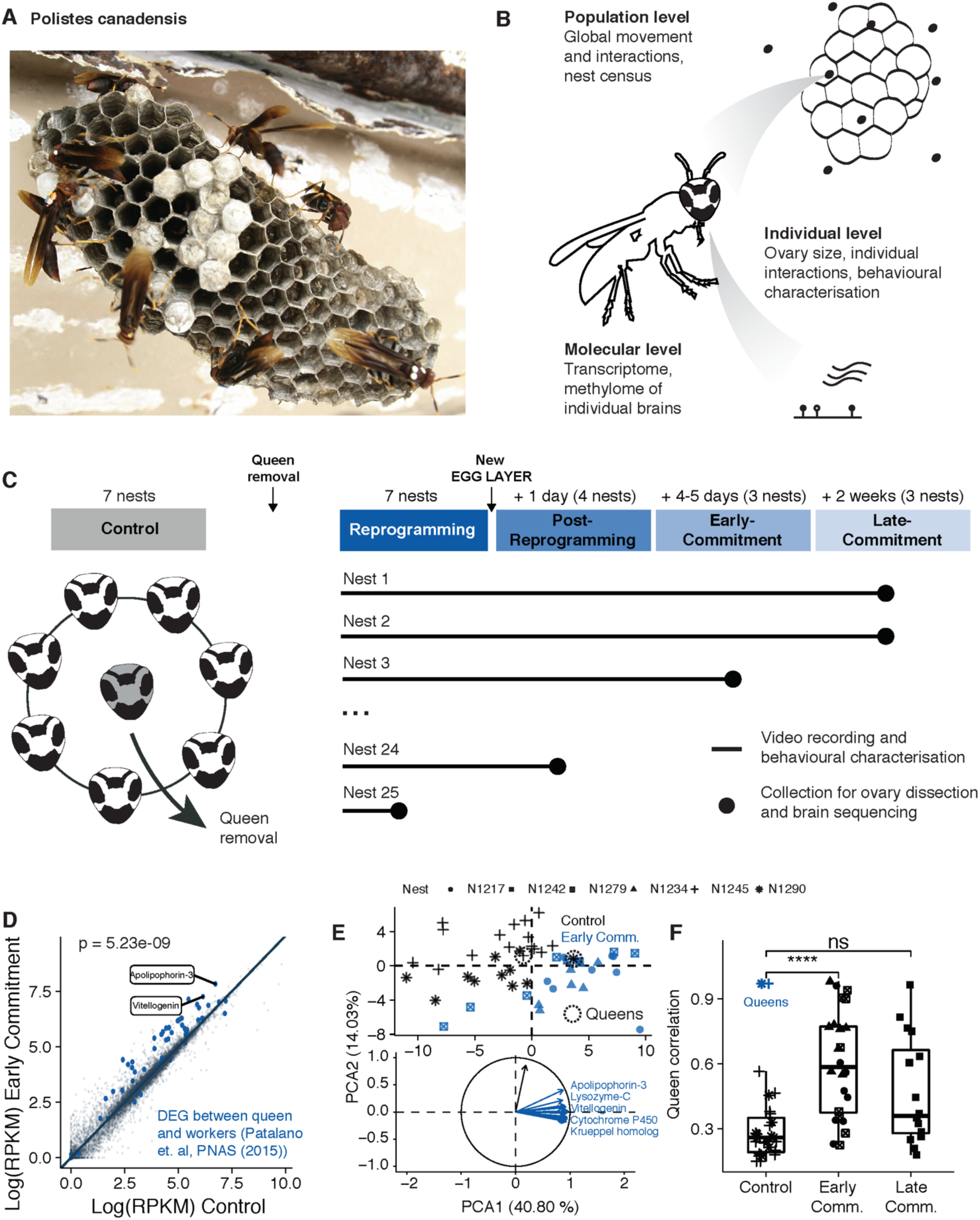
Upregulation of queen genes during reprogramming. **(A)** Photo of a typical nest of *Polistes canadensis* **(B)** Schematic of the multi-scale experimental approach **(C)** Schematic of the experimental time course of queen removal experiments. Control: 2-3 weeks prior queen removal; Reprogramming: time between queen removal and appearance of a new egg layer; Post-Reprogramming: Day and day+1 after appearance of a new egg layer; Early Commitment: day+4 or +5 after egg laying **(D)** Expression levels of genes (points) in the control against the early commitment phase. Up-regulated queen genes identified in (*15*) are marked in blue **(E)** Principal component analysis of individual insects (points) using queen gene expression levels. Only the 20 most contributor genes are shown in the radar plot of the variable of the PC **(F)** Correlation of the queen gene expression of all individual wasp from the experimental nests (points) compared to the queen gene expression of the queens in the control nests (in blue).

## Results

### Upregulation of queen genes in all workers after queen removal

Out of the 25 nests monitored in their natural environment, queens were removed from 18 nests 2 to 3 weeks after the first emergence of workers (queen’s daughters). After queen removal, a fast reprogramming phase begins, at the end of which (typically 6 days) a single new reproductive individual (queen) is established as evidenced by observation of egg laying (Fig. S1A). To understand the transcriptional changes associated with queen replacement, we performed brain RNA sequencing of all workers from 2 of the control nests (27 individuals) where the queen had not been removed and from 3 other nests on which the queen had been removed; these manipulated nests were collected 4 to 5 days after a new egg layer was detected (early commitment, 24 individuals). Workers in reprogrammed nests strongly up-regulated genes previously associated with reproductive phenotypes, henceforth referred to as “Queen Genes” (*13, 15*), and particularly *Vitellogenin* or *Apolipophorin-3* involved in lipid transport (Fig. 1D)(*18, 19*). A principal component analysis shows that the remaining workers in the early commitment phase obtain similar transcriptomic profiles largely due to queen gene expression (Fig. 1E). This result was further corroborated by these individuals displaying greater transcriptomic similarity with the control queens than with control workers, showing a transient loss of the worker phenotype during reprogramming which is eventually restored two weeks later upon emergence of an established queen (late commitment nest,16 individuals) (Fig. 1F).

### Increase of dominance behavioural interactions during reprogramming

Although this convergence of molecular identities towards the missing phenotype is presumably necessary for the eventual appearance of a new queen, it does not explain how the steady state with a single queen per nest is restored. We therefore reasoned that to obtain a single queen there needs to be a collective process which breaks the symmetry of molecular queen signatures between individuals on the population scale. To study whether there is a population-level component in the regulation of reprogramming and phenotypic specialisation, we analysed the recordings of 17 hours of videos from 5 colonies and quantified both global activity (overall movement at the nest level) and the individual interactions, such as dominance behaviours (Supplementary theory, Figs. 2A, S2A-B). We found an increase in both nest activity and the rate of interactions, particularly those involving fighting, during the reprogramming process suggesting that such interactions could contribute to the re-establishment of the steady state (Figs. 2B, S2C-D).

**Fig. 2.**
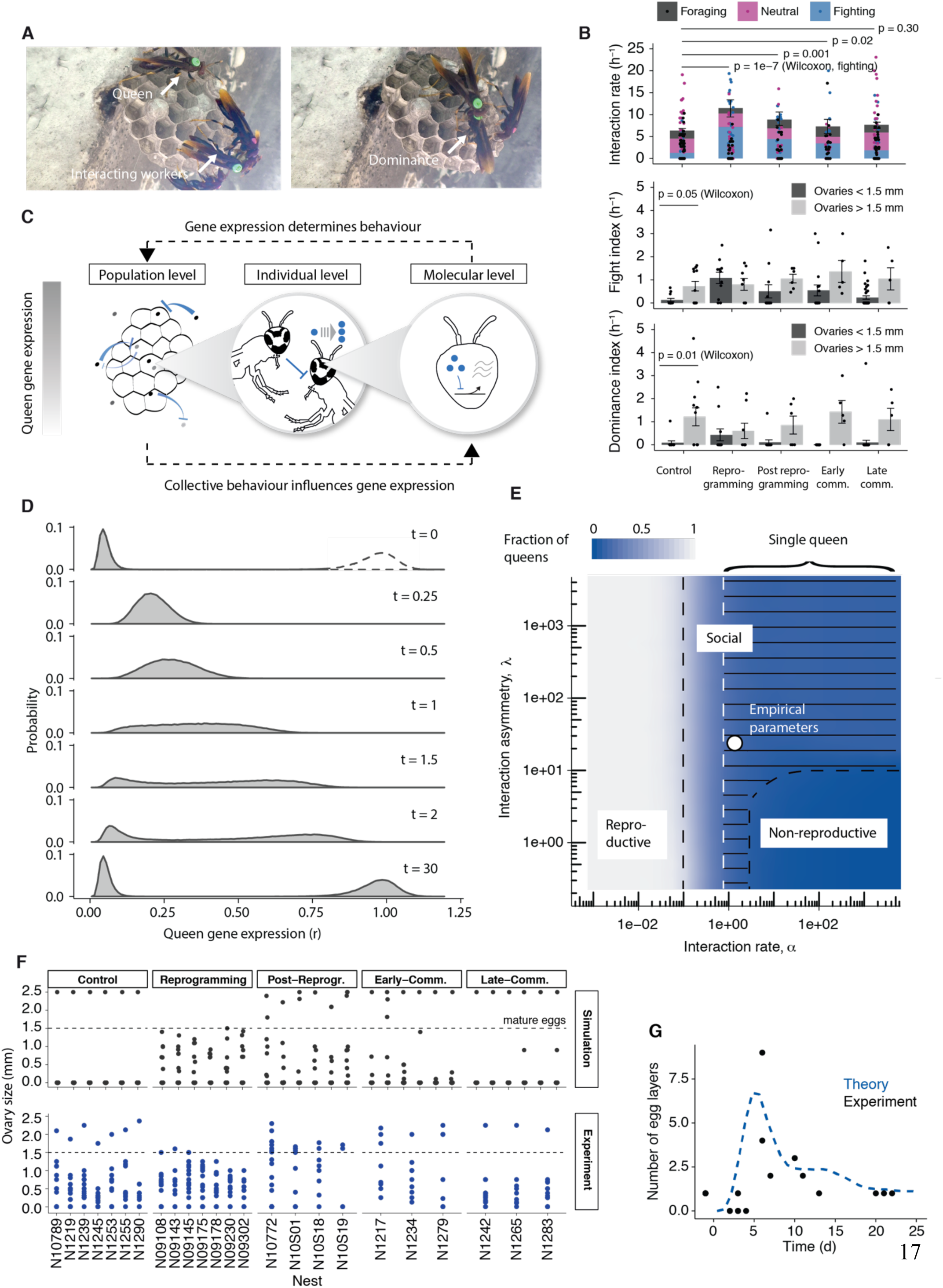
Plasticity and specialization emerge from antagonistic interactions on the molecular and the population scales. **(A)** Snapshots of videos showing the interaction between wasps **(B)** Quantification of the interaction rate separated by type (top), the fight index (middle) and the dominance index (bottom). Bars depict mean ± SEM and dots represent individuals. **(C)** Schematic of the model (more details in Supplemental Theory) **(D)** Probability densities of queen gene expression levels in a nest after queen removal obtained by stochastic simulations **(E)** Phase diagram showing different types of social structures as a function of the interaction asymmetry (*λ*) and the interaction rate (*α*). Empirical parameter values are marked by white circle **(F)** Experimental measurements of the size of the most mature egg of individual wasps for different nests across the whole duration of the reprogramming process (top). Exemplary stochastic trajectories (bottom) **(G)** Predicted and measured number of egg layers (ovary size > 1.5mm) over time after queen removal.

For each individual, we then calculated indices representing the frequency of their involvement in aggressive interactions (fight index) and the probability of dominance during these altercations (dominancy index), and classified individuals according to ovary size which can be used as a read out of queen gene expression (Sup. text, Fig. S2E). In control nests there is a strong asymmetry of interactions between individuals where only individuals with evidence of ovary development are involved in fights and successfully dominate nestmates (Fig 2B). This asymmetry broke down during the reprogramming phase where most individuals interacted in an aggressive manner consistent with the up-regulation of their queen genes as observed previously (*20*). Therefore, in order to break the molecular symmetry across wasps, fighting interactions must have a repressive effect on the expression of queen genes in the subdominant animal.

### Antagonistic dynamics on the molecular and population level allow the emergence of a single queen

To test the hypothesis that phenotypic specialisation results from the combination of molecular upregulation of queen genes and their population-level repression we used biophysical modelling (Fig. 2C). In this model, in the absence of behavioural interactions with the queen, individuals constitutively express queen genes producing and degrading the corresponding proteins(*21, 22*). In the context of a nest, behavioural interactions lead to the production of factors that repress the expression of queen genes in subdominant individuals (cf. Figs. 2B)(*23*). Therefore, in our model, collective dynamics on the population level influence queen gene expression and, vice versa, queen gene expression determines the rate and outcomes of interactions between pairs of individuals as in Fig. 2C.

Our calculations and stochastic simulations showed that, starting from a configuration of individuals with low expression levels of queen genes (workers) the competition between upregulation on the molecular scale and repression on the population level indeed leads to a collective upregulation of queen genes in workers after queen removal, ultimately giving rise to a sharply peaking bimodal distribution of phenotypes in the population (Fig. 2D). While experimentally determined parameters indeed lead to the emergence of a single queen, the model captures a range of potential population structures depending on the interaction rate and the degree of the sensitivity of dominance behaviour (asymmetry). More specifically, we predict that for weak and symmetric interactions all wasps have reproductive phenotypes, while for sufficiently frequent and asymmetric interactions exactly a single queen phenotype is guaranteed to emerge without the need for tuning of parameters (Fig. 2E).

With the model so defined we set out to challenge the validity of its mechanistic foundations by asking whether it was capable of quantitatively predicting the relaxation dynamics after queen removal across different scales. Indeed, our model quantitatively predicts the time evolution of ovary sizes and nest activity during the reprogramming dynamics (Figs. 2F-G, S2F), including the non-trivial transient emergence of multiple egg-layers after the reprogramming process (Figs. 2F-G).

### Specialisation and plasticity are simultaneously achieved by distinguishing between intrinsic and extrinsic perturbations

To gain mechanistic insight into how robust specialisation and plasticity are simultaneously achieved in the wasp society, it is instructive to consider the co-evolution of the molecular dynamics - represented by the gene-expression level of queen genes, *r*, and the population composition, represented by the number of individuals that have gene expression level *r* at time *t, f*(*r, t*) (Fig. 3A). To this end, we computed the time evolution of a hypothetical individual with a given expression value of queen genes and the corresponding evolution of the population composition (Fig. 3B, left, Video S2). The co-evolution of both scales is represented by black arrows, with an exemplary trajectory highlighted in blue. Our analysis showed that the dynamics relax to a steady-state composition of the population (represented by a specific distribution of queen gene expression values, *f*). This steady state comprises two attractors of the microscopic dynamics, one corresponding to the worker phenotype and one corresponding to the queen phenotype. These two attractors give rise to a bimodal population composition. These specialised phenotypes are subject to different sources of perturbations: intrinsic perturbations, such as gene expression noise, and extrinsic perturbations, such as the removal of the queen. Intrinsic perturbations affect individuals independently such that the population composition remains unaffected. Such a perturbation leads to gene expression dynamic in the opposite direction of the perturbation, such that this perturbation is actively suppressed by the interplay between molecular and population scale dynamics (Fig. 3B, middle, horizontal displacements). Extrinsic perturbations affect both the population composition, *f*, and gene expression levels, *r*, poising the system at a “plastic” state where individuals have the capacity to become either worker or a queen (Fig. 3B, right, diagonal displacement). Mathematically, the system undergoes a saddle node bifurcation with the population structure, *f*, as a functional bifurcation parameter (Fig. 3C). Therefore, *Polistes* integrates antagonistic dynamics on different scales to distinguish between intrinsic, molecular perturbations (which are uncorrelated between individuals) and extrinsic, population-level perturbations (which are correlated between individuals), reacting stably to the former ones and plastically to the latter ones.

**Fig. 3:**
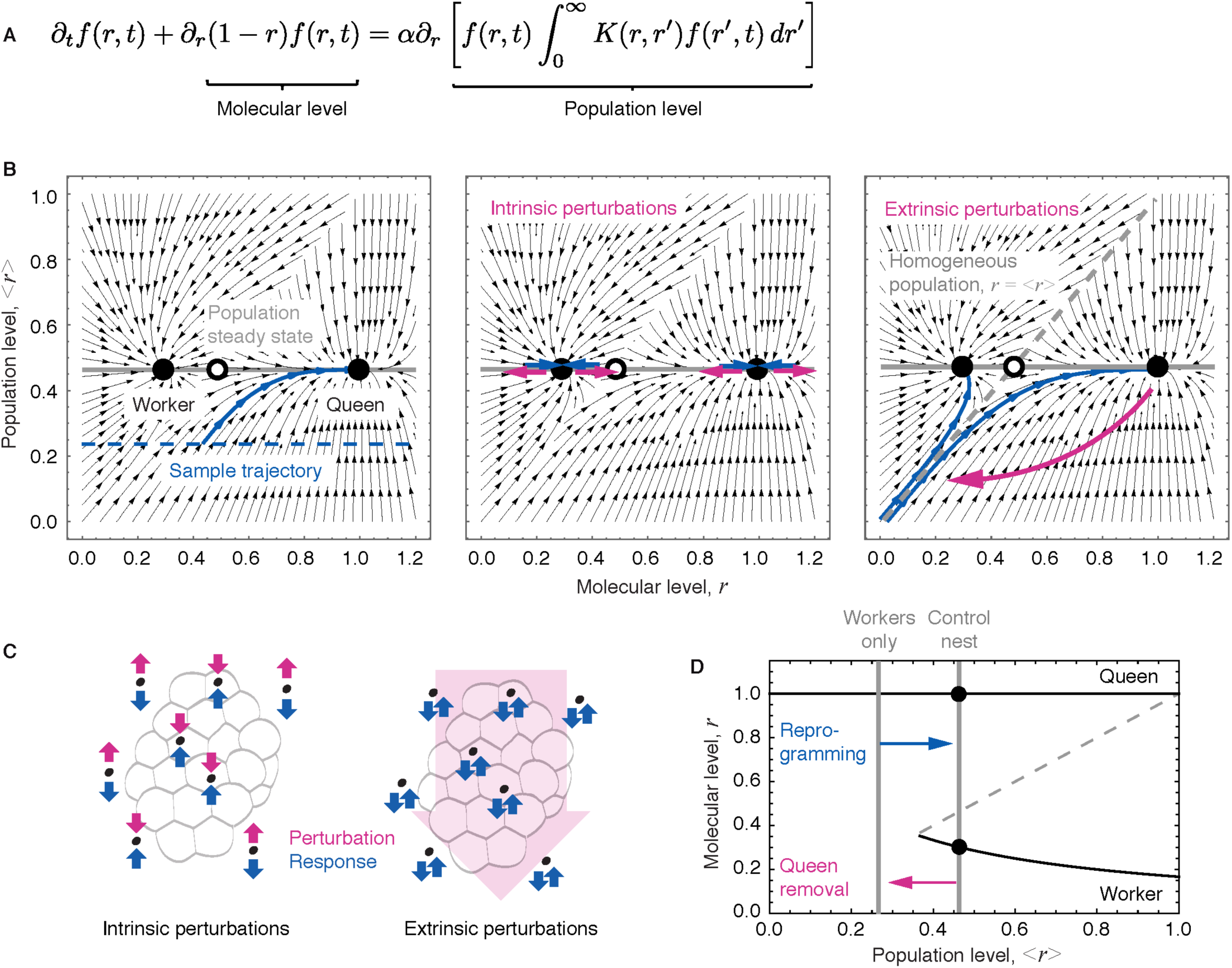
The colony reacts differently to perturbations across scales. **(A)** Mean-field master equation describing the time evolution of the number of insects *f*(*r, τ*) with queen gene expression level *r* at a given time *τ*. The interaction kernel, *K*(*r, r*′), defines the rate of interactions between individuals with queen gene expression *r* and *r*′ (Supplemental Theory). **(B)** Phase portrait depicting the joint dynamics of the population structure *f*(*r, τ*), represented by the average ⟨*r*⟩, and individual insect, represented by its queen gene expression, *r*. Left: unperturbed dynamics, middle: intrinsic perturbations, right: extrinsic perturbations. **(C)** Schematic depicting the response to intrinsic and extrinsic perturbations. **(D)** Bifurcation diagram. Stable attractors are depicted as solid black lines, unstable states are represented by the dashed grey line.

### DNA methylation contributes to long term stability at the population level

Our analysis shows how the paper wasp simultaneously achieves robust specialisation and plasticity. But how can this specialisation be stable in the long term? A *Polistes* worker is, on average, subject to a subdominant interaction 3.7±1.6 (SEM) times per day (Fig. S3A). Interactions therefore occur roughly on a similar time scale compared to typical gene activation and protein degradation times(*22*), necessarily leading to the frequent chance activation of queen genes in workers due to the stochastic timing of interactions (Fig. S3B, Supplemental Theory).

Indeed, analytical calculations and Monte Carlo simulations show that while the model predicts the emergence of a single queen *on average*, it also predicts that due to strong fluctuations the queen is replaced within days, whereas the queen state is stable in paper wasp societies for several weeks. This result suggests that additional factors stabilise wasp societies against such fluctuations.

As in mammals, *Polistes* have additional, epigenetic, layers of gene regulation, including chemical modifications of the DNA, such as DNA methylation(*15, 24*). Unlike in mammals, however, DNA methylation in *Polistes* is almost exclusively found in gene bodies. We performed brain whole genome bisulfite sequencing of the same individuals used for transcriptomic analysis and collected during the early commitment phase, allowing us to correlate both measurements with each other (Fig. 4A). Correlating bisulfite sequencing to RNA sequencing we found that DNA methylation is associated with the suppression of gene expression noise in *Polistes.* Specifically, in addition to positively correlating with expression levels as observed previously(*25*), DNA methylation in gene bodies reduces gene expression variability independently of the phases as measured by excess variance across workers over technical noise (Figs. 4B, S3C-D), highlighting the stabilising role of DNA methylation on gene expression in a primitively social wasp(*26*). Our theoretical work in turn shows that such a reduction in gene expression variance due to DNA methylation exponentially decreases the replacement rate of the established queen (Fig. 4C, Supplemental Theory). Hence DNA methylation, contributes to stabilising the population as a whole.

**Fig. 4:**
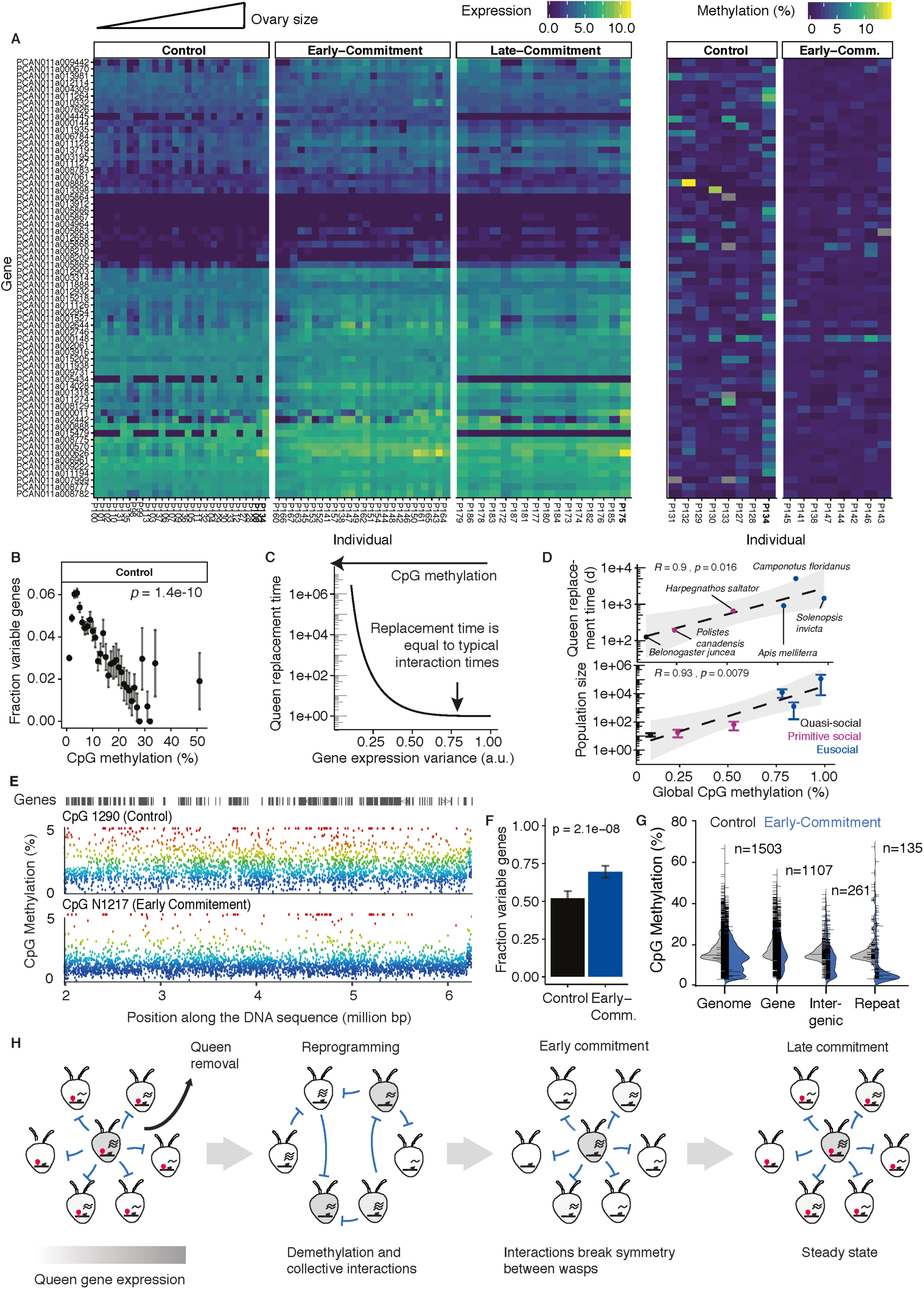
Epigenetic factors contribute to the stability of the social structure. **(A)** Heatmap depicting gene expression (left) and DNA methylation levels (right) of queen genes. Genes are ordered based on hierarchical clustering of gene expression in control nests. Individuals are ordered by their ovary size (small to large, left to right) and queens are marked in bold. **(B)** Fraction of genes showing significant biological variability between workers (see Material and Methods) binned by similar DNA methylation levels in control nests **(C)** Theoretical prediction of queen replacements times as a function of gene expression variance **(D)** Queen replacement times (top) and typical colony sizes (bottom) as a function of global CpG methylation different species (dots) and social organization (colour) **(E)** Example of global demethylation. Every dot represents average methylation across individuals in a window containing 50 informative CpGs. **(F)** Fraction of significantly variable genes in control and early-commitment phase **(G)** Distribution of CpG methylation levels for different genomic features **(H)** Graphical summary of the sequence of events during relaxation of the nest.

Consistent with this paradigm, we reasoned that the higher the DNA methylation level in the genome the more stable the social structure should be (Sup. Text). Drawing on quantitatively comparable methylation studies across different population structures of social insects species (*26*) together with new data generated in this study on normalised mass-spectrometry results obtained from the quasi-social wasp *Belanogaster juncea*(*27*), we found indeed that the typical replacement time of the queen phenotype strongly correlates with global DNA methylation levels (Figs. 4D-top S3E, Supl. Text). Further, the stabilising effect of DNA methylation should be more relevant in large colonies where the rate of subdominant interactions per individual is restricted. Although genetic, developmental and nutritional factors are also implicated in complex social (eusocial) organisation(*6, 28, 29*), we found that global levels of DNA methylation were positively correlated with typical colony size (Fig 4D-bottom). These results support the conserved role of DNA methylation in social organisation(*12, 30–32*)

### Erasure of DNA methylation during reprogramming

If DNA methylation stabilises phenotypic specialisation in the long term, we would expect methylation to be depleted during reprogramming. In mammalian cells reprogramming is associated with the erasure of DNA methylation marks(*33*). Indeed, we observed a partial erasure of DNA methylation in the early-commitment nests compared to stable control nests (Fig.4E). Our results obtained by bisulfite sequencing and validated by mass spectrometry showed that among the 5 nests collected during the post-reprogramming phase, all reprogrammed individuals had lower DNA methylation levels than control individuals (Figs. 4A, S4A). The change in DNA methylation levels of queen genes between control and early commitment nests follows the general trend of DNA methylation erasure (Fig. S4B). Accordingly, their gene expression variability increased during early-commitment (Fig 4G). While depletion of methylation is evident in all genomic features (Fig. 4F) we found 198 genes who either significantly acquired methylation or in which demethylation was more pronounced than the genome average, respectively (Fig. S4C). Functional enrichment analysis of these genes showed that they were mainly involved in protein transport and particularly in lipid transport, including the gene Nedd-4 previously described to have a role in remodelling neuronal circuitry (Fig. S4D-E)(*34*). Therefore, our results show that, similar to mammals, DNA methylation is a flexible epigenetic mark regulated by social environment and, potentially, among other molecular factors, able to feed back onto the stability of the social structure(*35, 36*). Future studies will be required to establish the molecular pathways governing dynamic DNA methylation in *Polistes*.

## Conclusions

Both experimental and theoretical approaches show that *Polistes* uses antagonistic dynamics on different spatial scales to distinguish between molecular and population-level perturbations, thereby achieving robustness to the former and plasticity to the latter. Our findings also suggest that epigenetic DNA modifications play an unanticipated functional role in regulating the stability of the society at the population level. Our work demonstrates that correlated measurements across scales can give qualitatively new insights into the mechanisms underlying self-organisation of biological systems (Fig. 4H). Our approach may be more widely applicable to other biological systems of interest.

## Acknowledgments

We thank Frank Jülicher, Benjamin D. Simons, and all members of the Reik laboratory for helpful discussions. We also thank Nathalie Smerdon at the Wellcome Trust Sanger Institute and Felix Krueger and Simon Andrews at the Babraham Institute for processing sequencing data and bioinformatics support with Illumina sequencing. We thank T. Lengronne, R. Zaurin, R. Southon, E. Bell for help in the field, all the staff at the Galeta field station and at the Smithsonian Tropical Research Institute Panama for help and logistical support in fieldwork. We also thank P. Vardakas for assistance in the video analyses. This work was conducted under under Autoridad Nacional del Ambiente (ANAM) permits #SE/A-33-09, #SE/A-65-10, #SE/A-20-12, and export permit 10BR004553/DF and 11BR006471/DF.

## Author contributions

A complete list of contributions to the paper is available in Table S1.

## Data and materials availability

All others data and material used in the manuscript is available in the main text or the supplementary materials. Source code is available upon request to the authors.

