## Supplementary Material for "Specialisation and plasticity in a primitively social insect"

**This PDF file includes:**

Materials and Methods
Supplementary Text
References
Supplementary Figure legends S1 to S4
Table S1
Captions for Movie S1
Supplementary Theory

**Other Supplementary Materials for this manuscript include the following:**

**Movie S1.** Computational vision of global nest movement analysis
**Movie S2.** Stochastic evolution overlaid on the phase portrait

#### Materials and Methods

Fieldwork and sample collection

*Polistes canadensis*: Field experiments were conducted between June and August during 3 expeditions to Panama between 2009 and 2012 in the protected area of Punta Galeta, Colon (9°21'30.877''N, 79°54'0.0053''W), under the field collection permits #SE/A-33-09, #SE/A-65-10 and #SE/A-20-12 from the Autoridad Nacional del Ambiente (ANAM) of Panama. Nests were all selected at pre-emergence stages, when only the queen and few co-foundresses were present. All nests were monitored daily in order to mark every new emergent worker and censused every other evening to know the entire nest population. The queen was identified by removing an egg and observing who replaced it, and also from behavioural observations and census data. Nest were monitored for three to nine weeks and then female wasps representing the different stages of the reprogramming and commitment phases that take place during queen succession were collected. Phenotypic reprogramming was induced by removing the queen and an egg. Nests were monitored every day to detect any new egg layers. As soon as a new egg appeared the reprogramming phase (where a worker transitions to be a queen) was considered to be over, as a new egglayer had been established. The identity of this new egg layer was determined over the following 2 days by behavioural observations; this corresponds to the post-reprogramming phase. The wasps from 5 nests were collected at this stage. A further 6 nests were monitored further, generating samples of post-reprogramming wasps on nests 4 to 5 days after the first egg laying occurred (early commitment phase), at which point they were collected. A further 3 nests were monitored until 2 weeks after egglaying (late commitment phase) at which point they were collected. All collected nests have similar stage of development (35,6 ±14 days post-emergence) and comparable size (15,6 ± 7,2 individuals) and without any sign of parasites or disease. Wasps were collected directly off their nests individually with forceps during the active hours of the days. Their heads were cut off and immediately placed in RNA later solution (Ambion). Their bodies were stored in 80% ethanol and kept at -20°C until dissection of ovaries. Reproductive state was assessed for 378 wasps by measuring the first and biggest egg at the entrance of the oviduct. A mature egg has a size of 1.5 mm, which corresponds to the smallest egg size observed and associated with egg-laying.

*Belonogaster juncea*: Three nests of *B. juncea* and their 11 individuals were collected in March 2013 in Ebolowa, Cameroon (2°55′N 11°9′E). Collection, storage and reproductive state assessment of all individuals were done similarly than for *P. canadensis*.

Liquid chromatography – Mass-spectrometry

Genomic DNA were isolated from 64 single brains of wasp using protocol from (*1*). For the detection of methylgroups, five hundred nanograms of genomic DNA was incubated with 5 units of DNA Degradase Plus (Zymo Research) at 37 °C for 3 h. The resulting mixture of 2-deoxy-nucleosides was analyzed on a Triple Quad 6500 mass spectrometer (AB Sciex) fitted with an Infinity 1290 LC system (Agilent) and an Acquity UPLC HSS T3 column (Waters), using a gradient of water and acetonitrile with 0.1% formic acid. External calibration was performed using synthetic standards, and for accurate quantification, all samples and standards were spiked with isotopically labeled nucleosides(*2*).

RNA-Sequencing

Total RNA was extracted from 87 single brains, using theQIAGENAll PrepDNA/RNAMini Kit according to the manufacturer's instructions. 50 to 200 ng of total RNA was enriched for mRNA using Dynabeads Oligo(dT)25 from Invitrogen in two subsequent steps of purification with fresh beads. Fragmentation was done by incubation of mRNAs for 5 min at 94 °C in the First-Strand Buffer (Invitrogen), and directly followed by cDNA synthesis using a SuperScript III Kit (Invitrogen) according to the manufacturer's instructions. dUTPs were incorporated for second-strand synthesis for library orientation except for samples P188 to P212. cDNA was end-repaired, A-tailed, and ligated with a methylated Adaptor Oligo Kit (Illumina) using the NEB Next Kit (New England Biolabs) according to the manufacturer's instructions. dUTP excision was done before amplification using USER mix (New England Biolabs). Libraries were amplified with 16 cycles using 2× Phusion HF buffer (New England Biolabs). Size selection and cleaning between steps were performed with the AMPure XP system (Agencourt) to select DNA fragments between 250 bp and 500 bp. Paired-end libraries were sequenced on HiSeq 2500 Illumina platform.

Whole-Genome Bisulfite Sequencing

Genomic DNA was extracted from 16 single brains, using the QIAGEN All Prep DNA/RNA Mini

Kit according to the manufacturer's instructions. Between 200 ng and 500 ng of input genomic DNA was used per library and spiked with unmethylated lambda DNA to provide an estimation of BS conversion efficiency. DNA was end-repaired, A-tailed, and ligated with a methylation Adaptor Oligo Kit (Illumina) using the NEB Next kit according to the manufacturer's instructions. The adaptor-ligated DNA was treated with sodium-BS using an Imprint DNA Modification Kit from Sigma–Aldrich according to the manufacturer's instructions for the two-step protocol. BS-treated DNA was amplified using KAPA HiFi Uracil + DNA Polymerase (KAPA Biosystems) with 15 cycles. Size selection and cleaning between steps were performed with an AMPure XP system (Agencourt) to select DNA fragments between 250 bp and 500 bp and sequenced at the Sanger Institute using the HiSeq 2500 Illumina platform.

RNA sequencing analysis

Libraries were sequenced on the Illumina HiSeq platform using the default RTA analysis software. RNA-Seq data data were trimmed with Trim Galore (v0.4.1, default parameters) and mapped to the *Polistes canadensis* genome assembly GCF_001313835.1 using TopHat v2.0.12 as previously described in (*1*). Strand specific quantification was performed using RNA-seq pipeline in Seqmonk software Version 1.39.0 (www.bioinformatics.babraham.ac.uk/projects/seqmonk/). Gene expression levels were normalized in terms of reads per million (reads per kilobase per million, RPKM). To normalize across nests a scaling factor was calculated using the DEseq2 package in R and log transform with only orientated samples. "Queen genes" list correspond to Queen-biased genes identified in (*1*). PCA analysis was performed using FactoMineR package in R. Genes contributing the most to a given PCA dimension were identified from their cos2. Hierarchical cluster analysis was performed using heatmap.2 package in R. Statistical analyses were performed using R. To calculate biological variance, after removing control queens from the dataset we fitted a technical noise model using the trendVar function of the scran package (version 1.12.1) and computed noise contributions using scran's decomposeVar function. Genes with p-values smaller than 0.1 after multiple testing correction using the Benjamini-Hochberg method were considered significantly variable.

Bisulfite sequencing analysis

Libraries were sequenced on the Illumina HiSeq platform using the default RTA analysis software.Raw sequence were trimmed to remove both poor quality calls and adapters using Trim Galore (version 0.3.5 with default parameters, www.bioinformatics.babraham.ac.uk/projects/trim_galore/). The remaining sequences were then aligned to genome assembly (EVM/PASA; Patalano et al. PNAS, 2015) using Bismark (version 0.12.2, with the parameters: --bowtie2 --score_min L,0,−0.4) and Genome_build: Polistes canadensis GCF_001313835.1 from (1). Quantification was done in SeqMonk over probes which contain 50 CpGs each with a minimum read count of 4. Only 30pb to 1kbp probes were kept. Intergenic and gene features analysis was done using probes that specifically overlap these features. Repetitive genomic regions were identified using RepeatMasker v4.0.1, using the 20120418 repeat libraries limited to Apocrita species. Methylation over a given feature was calculated by averaging the methylation levels of CpGs across the probes overlapping a given feature. Binomial filter from Seqmonk was applied to identify methylated probes having statistically greater variations than the ones observed globally (using CpGs covered by a minimum of 4 reads). Significance was considered only when probes deviate from the global profile by at least 10% after Benjamini-Hochberg corrections. GO enrichment analysis was perform with OmicsBox using Fisher Exact test ($p<0.01$) and reduce to the most specific terms.

Video Analysis

A HD Video Camera was placed in front of the nests during the active hours. We analysed over 17hours of videos from 5 colonies from 2012 during the 5 phases (average of 50 min/phase/nest). For the global movement analysis of the overall nest, camera movements and unusual captures that might disrupt the quantification were removed. Computer vision analyses were performed by the quantification of the global pixel changes measured between each frame using Python programming language and Open CV (Open Source Computer Vision Library) and normalized by the number of wasps' present. To allow phase and nest inter-comparisons, a normalization of a nest distance with the camera was computed by measuring the diameter of the label used to mark the wasps (Fig. S2A).

For the individual behavioral analyses, we monitored a total of 821 interactions from 71 individuals (average of 87min/wasp) and classified them into 6 different types of interactions as previously described in *Polistes* (*3*, *4*). Briefly, DOM: domination over a nestmate (bite, sting, chew, nip or chase). SUB: subordination (in response to DOM or escape). DEC: donor of foraging material to a nestmate. REC: receiver of the DEC. TR: involved in trophallactic exchanges (i.e. exchanges of fluid from one wasp to another). ANT: Involved in antennation. For Fig. 2B we categorised these types as follows: Fighting as DOM and SUB, foraging as REC and DEC, and neutral interactions as ANT and TR. From this table, Interaction rate (DOM+SUB+DEC+REC+ANT+TR per hour per wasp), Fight index (DOM + SUB / All interactions) and DOM index (DOM/(DOM+SUB)) were calculated for each individual wasp and normalised by the amount of time each wasp was observed on the nest. Wasps present in the nest but not performing any interaction were also included in our analysis. To calculate the rate of subdominant interactions per worker and per hour, i.e. a passive measure of interactions, we took into account those individuals who were not present in the nest during the video in order to obtain an unbiased measure of this rate. In this specific case, we therefore normalised by the total number of individuals detected in a night-time population census conducted before the video recording.

#### Supplementary Text

Ovaries size as a read out of queen-biased gene expression

It has previously been reported for *Polistes* and other species that ovarian development is mediated through Juvenile Hormone (JH) and activation of the *vitellogenin* gene in individuals engaged in successful fighting interactions(*5–7*). Therefore, ovary size is used in Fig. 2B as a proxy for queen gene expression (*vitellogenin* and *apolipophorin-3*).

Methylation survey on different types of social insect organization

An analysis of several social hymenoptera predicted a direct link between the methylation rate of their genomes and the degree of complexity of their societies (based on the number of individuals and the castes that compose it in stable societies)(*8*). Despite the increasing number of sequenced methylomes in social insects, comparative studies are still rare and mainly contradictive due to the discrepancy of the methodologies used to analyze sequencing products. Therefore, we took advantage of a recent comparative publication of five social hymenopterans, which re-quantified the percentage of methylation since the original reads, and associated them with two measures for complexity and stability of societies (*9*). The methylation levels of the quasi-social organization of *Belonogaster juncea* wasps, where more than one individual in a nest have sign of developed ovaries, were included in this analysis (*10*). We measured their global level of brain DNA methylation by Mass-Spectrometry and normalization of their overall levels was performed through those of *Polistes canadensis* present in the two studies. We show that both number of individuals and queen turnover are positively and significantly correlated with the degree of complexity of social organizations (Fig. 4D), thus confirming previous predictions.

#### Supplementary Figure legends

**Fig. S1.**

**(A)** Reprogramming times (i.e. the time taken for a new egglayer to emerge) after queen removal for individual nests (dots). Mean and standard deviation are represented by crossbar.

**Fig. S2.**

**(A)** Normalisation using the tag diameter to adjust camera and nest distance **(B)** Example of an increase in global movement detection in the nest N1217 show by the increase of the amount of white at different time points of the analysis of pixel changes across various frames. 'Frame' indicates the time in which the picture was taken. 'Mov' indicates the number of pixels that changed compared with the previous frame **(C)** Global activity for each nest over time **(D)** Global changes in nest activity. Each grey dot represents a nest. Black dot and error bar signify mean and SEM of all frames. **(E)** Correlation between ovaries development and both *Vitellogenin* and *Apolipophorin-3* genes. Examples of specific ovary dissections for each phase are shown below **(F)** Prediction of global changes by the model.

**Fig. S3.**

**(A)** Rate of subdominant interactions per worker and per hour and normalised by the total number of individuals detected during night census (see Sup. Text) (B) Sample trajectory obtained from numerical simulations showing the stochastic turnover of queens (C) Mean gene expression as a function of the DNA methylation levels in gene bodies (D) Fraction of significantly variable genes as a function of DNA methylation in the same genes in the early-commitment phase (E) Global level of DNA methylation measured by mass spectrometry in *Polistes canadensis* and *Belonogaster juncea* control nests.

**Fig. S4.**

**(A)** Global level of DNA methylation measured by mass spectrometry across 5 nests expressed as a percentage of all cytosines. Statistical test was performed between all individuals from control nests versus all individuals from reprogrammed nest (p= 0.0065, t-test) **(B)** Log2 fold change of DNA methylation as a function of methylation in control phase **(C)** Scatter plot showing DNA methylation levels of unbiased genomic probes in control and early-commitment nests. Statistically significant (binomial test) enriched or depleted probes are marked in blue or red, respectively **(D)** Functional enrichment analysis of statistically significant enriched probes overlapping with genes **(E)** CpG methylation across Nedd-4 gene for 8 individuals from control nest (blue tracks) and 8 individuals from early-commitment nest (pink tracks). CpG methylation was quantified at base pair level for visualisation. Scale ranges from 0% methylated to 35% methylated (highest bar). Top annotation tracks show Nedd-4 gene structure (blue), the 30 to 1kpb CpG windows used for quantification (empty black boxes) and significantly variable window between control and early-commitment nests (Window and cluster are highlighted in pink).

**Table S1. Author contribution**

| Role | Name | Definition |
|---|---|---|
| Conceptualization | S.P., A.A., S.S, C. G-R., W.R., S.R. | Ideas; formulation or evolution of overarching research goals and aims. |
| Data curation | S.P., A.A., I.G., C.G-R., S.R. | Management activities to annotate (produce metadata), scrub data and maintain research data (including software code, where it is necessary for interpreting the data itself) for initial use and later re-use. |
| Formal analysis | S.P., A.A., C.G-R., S.R. | Application of statistical, mathematical, computational, or other formal techniques to analyse or synthesize study data. |
| Funding acquisition | S.S., W.R., S.R. | Acquisition of the financial support for the project leading to this publication. |
| Investigation | S.P., P.N., S.D. | Conducting a research and investigation process, specifically performing the experiments, or data/evidence collection. |
| Methodology | A.A., S.R. | Development or design of methodology; creation of models. |
| Project administration | S.P, W.R, S.R. | Management and coordination responsibility for the research activity planning and execution. |
| Resources | S.S., W.R., S.R. | Provision of study materials, reagents, materials, patients, laboratory samples, animals, instrumentation, computing resources, or other analysis tools. |
| Software | S.P., A.A., C.G-R., S.R. | Programming, software development; designing computer programs; implementation of the computer code and supporting algorithms; testing of existing code components. |
| Supervision | S.B, S.S., W.R., S.R. | Oversight and leadership responsibility for the research activity planning and execution, including mentorship external to the core team. |
| Validation | M.B | Verification, whether as a part of the activity or separate, of the overall replication/reproducibility of results/experiments and other research outputs. |
| Visualization | S.P., A.A., S.R. | Preparation, creation and/or presentation of the published work, specifically visualization/data presentation. |
| Writing – original draft | S.P., A.A., W.R., S.R. | Preparation, creation and/or presentation of the published work, specifically writing the initial draft (including substantive translation). |
| Writing – review & editing | | Preparation, creation and/or presentation of the published work by those from the original research group, specifically critical review, commentary or revision – including pre- or post-publication stages. |

**Additional data files**

**Video S1. Computer vision analysis of global nest movement.** This video shows an extract from the global movement analysis performed by computer vision. Each darkened pixel corresponds to a change compared to the previous frame.

**Video S2. Stochastic evolution overlaid on the phase portrait.** The video shows the results of a stochastic simulation of the dynamics of a nest consisting of 10 individuals (red dots) who are initially prepared in a worker state. Black arrows depict the flow of the mean-field dynamics (Supplemental Theory).

### Supplemental Theory

Adolfo Alsina and Steffen Rulands

February 15, 2020

In this Supplemental Theory we provide details of the derivation of the mathematical framework underlying the results presented in the main text.

### Contents


**1 Introduction** **3**

**2 Model definition** **5**

2.1 Dynamics on the molecular level . . . . . . . . . . . . . . . . . . . . . 5

2.2 Interactions between individuals . . . . . . . . . . . . . . . . . . . . . 6

2.3 Coupling between population-level interactions and gene expression . . 8

**3 Phase diagram** **10**

**4 Derivation of the mean-field master equation** **12**

4.1 Continuum limit . . . . . . . . . . . . . . . . . . . . . . . . . . . . . 15

**5 Derivation of the phase portrait** **16**

**6 Stability of the steady state** **22**

**7 Prediction of experimental data** **26**

7.1 Marginal distribution of ovary sizes . . . . . . . . . . . . . . . . . . . . 26

7.2 Experimental parameters . . . . . . . . . . . . . . . . . . . . . . . . . 29

**8 Numerical simulations** **31**

### 1 Introduction

The primitive social insect *Polistes canadensis* forms societies composed of a single queen, which is the only reproductive individual, and multiple workers. After removal of the queen the remaining workers are capable of reprogramming and producing a new queen. In this study, we therefore use *Polistes canadensis* as a model system to understand how biological systems form stable structures in noisy environments which can, at the same time, be rapidly remodelled upon specific cues.

In mathematical terms, within the scope of this work, we define *specialisation* to denote situations where the distribution of phenotypes in the steady state of the population exhibits clearly separated modes which are stable on time scales much longer than the intrinsic time scales of the system. Such a scenario is typically the result of effective barriers resulting either explicitly from a predefined potential landscape or implicitly, as entropic barriers, from the kinetic rules governing the dynamics.

Given such a situation, one would naively expect that the reestablishment of the population structure after removal of one of the modes occurs on similar time scales as the lifetime of the metastable steady state (*1*). With the term *plasticity* we denote the capability of the population to repopulate a missing mode on a time scale similar to the intrinsic time scales, i.e. much faster than the lifetime of the metastable steady state.

Our approach to understanding specialization and plasticity in the *Polistes* society combines multi-scale experimental measurements with biophysical modelling. Different nests of *Polistes* were monitored in video, their individuals dissected and their brains subjected to multi-modal sequencing (see details in Supplementary Material), both before and at different time points after queen removal. This experimental approach provides information on different levels of biological organisation: the social, individual and molecular scales. Integrating the information provided by these experiments on different scales allows us to understand the mechanistic principles underlying the robust specialisation and rapid reprogramming in *Polistes* societies.

In this supplement, we provide details of the calculations underlying the derivation and analysis of a multi-scale biophysical model to understand the simultaneous capability for robust specialization and plasticity in the *Polistes* society. In our approach we seek to define the simplest model that capable of describing the experimental phenomenology. Specifically, we do not assume non-linearities unless motivated by experimental observations. Although our model therefore necessarily does not reflect the full complexity of wasp behaviour, such a reductionist approach will allow us to obtain mechanistic understanding underlying the principles governing specialisation and plasticity. Starting from a model that describes the non-Markovian dynamics of the joint probability distribution we derive the time evolution equations for the marginal probabilities of gene expression levels and ovary sizes. We then take the mean-field limit to derive the structure of the phase space.

The structure of this document is as follows: in section 2 using a master equation formalism we construct a model that describes molecular processes taking place in each individual which we then couple through interactions on the population scale. In section 7 we draw on this model to derive the time evolution of experimental observables, such as the distribution of ovary sizes and global nest activity. Then, in section 3, we construct the rich phase diagram as a function of the interaction rate and the interaction sensitivity. In section 4 we derive a mean-field master equation from the full stochastic model describing the time evolution of the individual and collective dynamics. These results are then used in section 5 to build a phase portrait of the system, which we use to understand how specialization and plasticity are regulated. In section 6 we consider the stability of the systems with respect to fluctuations and investigate the stabilising effect of epigenetic processes on the social structure. Finally, in section 8 we describe the numerical implementation of the model.

### 2 Model definition

#### 2.1 Dynamics on the molecular level

Our RNA-seq analysis showed that genes, which are upregulated in queens compared to workers in control nests (queen genes), are collectively expressed in all workers during the reprogramming process. This suggests, on the one hand, that the expression of these queen genes can be described by a single degree of freedom. We here denote this degree of freedom on the molecular level by $n_i$, taken to be the total abundance of proteins corresponding to queen genes in individual $i$. On the other hand, our analysis demonstrates that these queen genes are constitutively expressed in the absence of queen interactions, a finding corroborated by previous studies on individual workers in Polistes and other social insects (*2*). Therefore, in the absence of interactions, the dynamics on the molecular level are described by the production and degradation of proteins with rates $\mu$ and $\delta$, respectively. The time evolution of the probability of finding protein abundances $\{n_i\}$ in a population of $N+1$ insects, $P(\{n_i\}, t)$, is governed by a master equation of the form

$$\frac{d}{dt}P(\{n_i\}, t) = \sum_{i=1}^{N+1} \mu \left[P(\{n_i - 1\}, t) - P(\{n_i\}, t)\right] + \delta \left[(n_i + 1)P(\{n_i + 1\}, t) - n_i P(\{n_i\}, t)\right] \,, \tag{1}$$

where the first two terms describe the Poissonian production of proteins and the remaining terms their degradation. From here on, we will omit the time dependence of $P(\{n_i\}, t)$ unless necessary. The ensuing stochastic dynamics give rise to a steady state characterised by a distribution of protein abundances with an average value of $\mu/\delta$. Therefore, in the absence of interactions, the dynamics converge to a steady state where all individuals express queen genes.

#### 2.2 Interactions between individuals

To break the symmetry between individuals and select a single queen the dynamics on the molecular level need to be coupled to a collective process on the population level. In the context of a nest, individuals interact in different ways, including fighting. These fighting interactions are directed, meaning that in every interaction there is a dominant individual, the attacker, and a subdominant one.

The rate with which individual $i$ is subject to a subdominant interaction with individual $j$, $K_{ij}$, can be written as the total interaction rate between both, $a_{ij}$, times the conditional probability $b_{ij}$ that individual $i$ is subdominant in such an interaction, $K_{ij} = a_{ij}b_{ij}$. To calculate $a_{ij}$ we note that according to our video recordings, which we correlated with ovary size measurements in individual insects, the interaction rate of an individual insect increases with ovary size and the expression level of queen genes, $a_i \equiv \sum_{j \neq i} a_{ij} \propto n_i$. The pairwise interaction rate therefore is proportional to the probability that both individuals interact in a given time interval, $a_i a_j$, times the probability that this interaction involves individuals $i$ and $j$, $2/[N(N-1)]$, $a_{ij} = 2a_i a_j/[N(N-1)] \propto 2n_i n_j/[N(N-1)]$, such that we set $a_{ij} = \omega n_i n_j$ where $\omega$ is proportional to $2/[N(N-1)]$. To derive the conditional probability that individual $i$ is subdominant, $b_{ij}$, we resort to previous work showing that the outcome of an interaction is strongly determined by the concentrations of Insect Juvenile Hormone (JH), a hormone involved in ovaries development (*3*). Indeed, in our video recording of control nests we found that, in an interaction between two individuals the subdominant one is with high probability the one with the smaller ovaries. Given that ovary size is a proxy for queen gene expression, we take the conditional probability of an individual being subdominant to depend on the gene expression difference between the two interacting individuals, $b_{ij} \propto n_j - n_i$.

Taken together, the conditional probability of $i$ being the subdominant individual in an interaction between $i$ and $j$ is

$$b(n_i, n_j) = \Theta(n_j - n_i)\,. \tag{2}$$

The total rate of subdominant interactions that individual $i$ receives then takes the form

$$\omega \sum_{j \neq i} K(n_i, n_j) P(\{n_j\})\,, \tag{3}$$

with the interaction kernel being defined by

$$K(n_i, n_j) = n_i n_j \Theta(n_j - n_i)\,. \tag{4}$$

In order to derive the above form of the interaction kernel we have assumed that individuals can accurately measure the gene expression levels of the insect they are interacting with. A more realistic approach is to take into account that sensing of gene expression levels is associated with an uncertainty. If this uncertainty is distributed following a normal distribution with zero mean and variance $\sigma^2$, the probability of individual $i$ being subdominant in an interaction with individual $j$ is

$$\int_{-\infty}^{\infty} \frac{1}{\sqrt{2\pi\sigma^2}} e^{\eta^2/2\sigma^2} \theta(n_j - n_i + \eta) d\eta = \frac{1}{2}\left[1 + \mathrm{Erf}\left(\frac{n_j - n_i}{\sqrt{2\sigma^2}}\right)\right]\,, \tag{5}$$

with the error function defined as $\mathrm{Erf}(x) = \int_0^x e^{-y^2}\, dy$. With this the interaction kernel reads

$$K(n_i, n_j) = \frac{n_i n_j}{2}\left[1 + \mathrm{Erf}\left(\frac{n_j - n_i}{\sqrt{2\sigma^2}}\right)\right]\,, \tag{6}$$

where $\sigma^2$ determines how precisely an individual can measure gene expression of other individuals. The above expression is difficult to manipulate both numerically and analytically due to the presence of the error function. It is, however, well approximated by another sigmoidal function

$$K(n_i, n_j) = n_i n_j \frac{e^{-\lambda(n_i - n_j)}}{1 + e^{-\lambda(n_i - n_j)}} , \tag{7}$$

with $\lambda = 2\sigma^{-1}$ denoting the sensing sensitivity.

#### 2.3 Coupling between population-level interactions and gene expression

How do interactions on the population scale get translated into changes on the molecular scale? An interaction leads to the transient increase in the concentration of factors influencing queen gene expression in the subdominant individual. While the precise nature of the pathways and molecular species involved in this process are not fully known in *Polistes*, to break the symmetry between insects and to obtain a single queen they must counteract the gene expression dynamics and thus can only be repressive. This reasoning is also supported by previous studies in *Polistes* showing that inhibition of fertility-linked compounds requires physical interactions (*4*). In the following, we therefore refer to these factors as *queen gene repressors*. Based on this, we make a minimal set of assumptions to describe the coupling between interactions and gene expression dynamics:

1. Interactions lead to the transient presence of queen gene repressors in the subdominant individual and
2. these factors, while present, inhibit the expression of queen genes.

Not knowing the precise nature of the molecular pathways triggered by an interaction we model these repressive factors as a binary variably, $q_i \in \{0, 1\}$, representing the absence or presence of repressive factors in individual $i$, respectively. In our model, queen gene repressors inhibit the expression of queen genes which mathematically translates to a vanishing rate of the production of queen gene products. Therefore, the production rate of queen gene products reads $\mu(1-q_i)$. The dynamics of the repressive factors themselves are comprised of two processes: queen gene repressors are activated in the subdominant individual upon an interaction ($q_i = 1$) and then persist for a time drawn from a distribution $\Gamma(t)$. If the activity of queen gene repressors involves sufficiently many steps $\Gamma$ will be approximately normally-distributed with a variance much smaller than the mean. We therefore set $\Gamma(t) \propto \delta(t_{\text{per}} - t)$, where $t_{\text{per}}$ is the typical persistence time of queen gene repressors.

With this in mind we can now mathematically define the time evolution of the joint probability $P(\{n_i, q_i\})$. To this end, let us consider a population of $N+1$ individuals, each with two degrees of freedom: the number of queen gene products, $n_i$, and the state of presence of repressive factors, $q_i$. Taken together, the stochastic dynamics is described by in terms of the non-Markovian master equation of the form

$$\begin{aligned}\frac{d}{dt}P(\{n_i, q_i\}) = \sum_{i=1}^{N+1} \Bigg\{ & \mu(1-q_i)\left[P(\{n_i - 1, q_i\}) - P(\{n_i, q_i\})\right] \\ & + \delta\left[(n_i+1)P(\{n_i+1, q_i\}) - n_i P(\{n_i, q_i\})\right] \\ & + \Gamma(t_i^{\text{int}})P(\{n_i, 1\})(1-2q_i) + \omega \sum_{j \neq i} K_{ij} P(\{n_i, 0\}) P(\{n_j, q_j\})(2q_i - 1) \Bigg\},\end{aligned} \tag{8}$$

where $t_i^{\text{int}}$ is a realisation of a stochastic process defined by the time elapsed since the last subdominant interaction of individual $i$ evaluated at time $t$. Eq. (8) accounts for the full stochastic dynamics of the population, encompassing gene expression dynamics and the dynamics of the repressive factors. Rescaling the number of gene products and time appropriately, we obtain the master equation in dimensionless form,

$$
\begin{aligned}
\frac{d}{d\tau}P(\{r_i,q_i\}) = \sum_{i=1}^{N+1}\Big\{ & (1-q_i)\left[P(\{r_i-\epsilon,q_i\})-P(\{r_i,q_i\})\right] \\
& + \left[(r_i+\epsilon)P(\{r_i+\epsilon,q_i\})-r_iP(\{r_i,q_i\})\right] \\
& + \Gamma(\tau_i^{\text{int}})P(\{r_i,1\})(1-2q_i) \\
& + \alpha\sum_{j\neq i}K(r_i,r_j)P(\{r_i,0\})P(\{r_j,q_j\})(2q_i-1)\Big\},
\end{aligned}
\tag{9}
$$

with a dimensionless coupling $\alpha = \omega\mu/\delta^2$, time $\tau = \mu\cdot t$, rescaled queen gene expression levels $r_i = \delta n_i/\mu$ and $\epsilon = \delta/\mu$.

#### 3 Phase diagram

In order to gain insight into the range of possible behaviours of the system we performed stochastic simulations of the dynamics defined by Eq. (9) as described in section 8. We scanned the phase space of the system as a function of two parameters that control the coupling between the molecular and population scales: the interaction rate, $\alpha$, and the sensitivity, $\lambda$. For a given combination of these parameters we sampled 50 independent trajectories of the stochastic nest dynamics defined in Eq. (9). We set the population size to $N = 10$ individuals and $\tau_{\text{per}} = 1$. We took averages over the the asymptotic ovary size (for a definition see section 7) at a time long after the steady state had been reached, $\tau = 60$. The results are shown in Figure 2E of the main text.

In this phase diagram, we identify three regimes based on the value of the asymptotic number of queens, defined here as those individuals whose gene expression value is larger or equal than 0.8, which corresponds to 80% of the steady state value. In the limit $\alpha \ll 1$ interactions occur on a much slower time scale than the molecular processes, effectively uncoupling the molecular and the population scales and leading to a steady state where all individuals obtain a queen phenotype. On the other hand, for $\alpha \gg 1$, the asymptotic composition of the society depends on the sensitivity, $\lambda$: If $\lambda$ is small, $\lambda \ll 1$, subdominant interactions affect all individuals with roughly equal probability. As a result, we observe that the asymptotic dynamics converges to a state represented solely by individuals with low expression of queen genes (workers).

On the other hand, if the scale defined by the sensitivity $\lambda$ is at least of equal order than the typical variability of queen gene expression values, i.e. individuals are able to distinguish phenotypically relevant changes in gene expression, $\lambda > 1$, interactions break the symmetry between individuals and a bimodal social steady state arises. While multiple queens can exist for precisely defined values of $\alpha$, exactly one single queen is guaranteed to emerge as long as the interaction rate exceeds a threshold value and individuals are capable of distinguishing "macroscopic" gene expression states. Therefore, by integrating antagonistic dynamics on different spatial scales, *Polistes* societies establish a single queen robustly for a large range of parameters, avoiding the need for fine-tuning of parameters that fluctuate in time and from nest to nest.

To locate the empirical parameter values in the phase diagram we estimated the parameters from the experimental data presented in Figure 2B of the main text, obtaining $\alpha_{\mathrm{exp}} \approx 1$. To estimate $\lambda$ we counted the number subdominant queen interactions. Out of 17 interactions involving queens in the control and late-commitment phases we observed 0 interactions, where the queen was subdominant. The maximum likelihood estimate for the error rate using the beta-distribution as a (conjugate) prior therefore is 1/17. Using the definition of the interaction kernel we find an analytical expression for the error rate,

$$2\int_{-\infty}^{0} d\Delta r(1+\exp(-\lambda\Delta r)^{-1} = 2\ln 2/\lambda$$

Solving for $\lambda$ we find that $\lambda \approx 24$. According to the phase diagram, these parameters indeed lead to the emergence of a single queen in support of the proposed paradigm.

### 4 Derivation of the mean-field master equation

The model defined in Eq. (9) predicts many of the features of the reprogramming process and properties of the steady state. But its high dimensionality and non-Markovianity render it unsuitable for analytical treatment. In order to understand how specialization and plasticity are simultaneously achieved in the *Polistes* society we start from Eq. (9) and develop a continuum, mean-field description. We will then employ this continuum description in section 5 to explore the structure of the phase space as a function of the individual and collective degrees of freedom.

To begin, we consider the time evolution of a single "tracer" individual embedded in a nest with a given composition $P\left(\{r_i, q_i\}_{i=1}^N\right)$ and study the evolution of the probability of finding the tracer in the state $(r, q)$ given the nest composition, $P(r, q) \equiv P(r, q|\{r_i, q_i\})$. In this approach we consider the nest as a "bath", which is not affected by the tracer and which determines the fluctuations of the individual tracer dynamics. The master equation for the time evolution of the queen gene expression level and repressor state of the tracer individual reads

$$\begin{aligned}\frac{d}{d\tau}P(r,q) &= (1-q)\left[P(r-\epsilon,q) - P(r,q)\right] \\ &\quad + \left[(r+\epsilon)P(r+\epsilon,q) - rP(r,q)\right] \\ &\quad + \Gamma(\tau_{int}^i)P(r,1)(1-2q) + \alpha\sum_{j=1}^{N} K(r,r_j)P(r,0)P(r_j,q_j)(2q-1)\,. \qquad (10)\end{aligned}$$

To obtain an equation for the marginal probability $P(r)$ we first need to integrate out the queen gene repressors state, $q$. To this end, we introduce a trajectory dependent time (cf. Figure 1) as

$$\tilde{\tau}(t) = \int_0^\tau d\tau' \prod_{i\in\mathcal{I}} f(\tau_i - \tau')\,, \qquad (11)$$

where $\{\tau_i\}_{i\in\mathcal{I}}$ is the set of times when an interaction, and hence a $q: 0 \to 1$ transition,

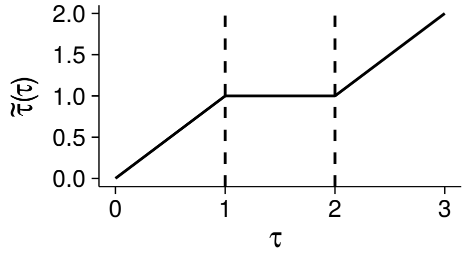


Figure 1: Example depicting the dependence of the trajectory dependent time $\tilde{\tau}$ as a function of the physical time. $\tilde{\tau}$ grows as the same rate as the physical time $\tau$ unless an interaction takes place (in this case at $\tau = 1$), in which case it remains at the same value for a time $\tau_{\text{per}}$.

takes place, and the function $f(\tau_i - \tau)$ is defined as

$$f(\tau_i - \tau) = \begin{cases} 0 & \tau \in [\tau_i, \tau_i + \tau_{per}] \\ 1 & \text{otherwise}\,. \end{cases} \tag{12}$$

From this definition it follows that $\tilde{\tau}$ increases in the same manner as $\tau$ if and only if $q = 0$ and is constant otherwise. The evolution of the system in the new time coordinate $\tilde{\tau}$ thus coincides with the evolution with respect to $\tau$ when $q = 0$ and its effect is collapsed to a single time point when $q = 1$. This is equivalent to considering dynamics where $q = 0$ for all times and where a suitably chosen number, $M$, of queen gene products are instantaneously degraded at times $\tau \in \{\tau_i\}_{i \in \mathcal{I}}$. $M$ is the typical number of molecules degraded in a time interval of length $\tau_{per}$, $M \approx r[1 - \exp(-\tau_{\text{per}})]$. The master equation describing such dynamics takes the form

$$\begin{aligned} \frac{d}{d\tilde{\tau}} P(r) = & \, [P(r-\epsilon) - P(r)] \\ & + [(\epsilon + r)P(r+\epsilon) - rP(r)] \\ & + \alpha \sum_{j=1}^{N} [K(r+M, r_j)P(r+M)P(r_j) - K(r, r_j)P(r)P(r_j)]\;. \end{aligned} \tag{13}$$

Next, we consider time scales much longer than typical interaction times. In this case, we consider time intervals, where the average number of degraded queen gene products is equal to $\epsilon$, of $M \to \epsilon$ while simultaneously rescaling the interaction rate $\alpha$ in such a way that the total number of queen gene products degraded in a long time interval, $\int_{\tau_1}^{\tau_2} d\bar{\tau}' M(\bar{\tau}_{per}) \prod_{i\in\mathcal{I}} \delta(\bar{\tau}_i - \bar{\tau}')$, remains invariant. Intuitively, such a coarse-graining operation corresponds to a homogeneous distribution of interaction events in the time domain for sufficiently long time scales. As the total effect of interactions on long time scales remains unchanged the structure of the phase portrait remains unchanged as well. With this, we obtain

$$\begin{aligned}\frac{d}{d\tilde{\tau}}P(r) = {} & [P(r-\epsilon) - P(r)] \\ & + [(r+\epsilon)P(r+\epsilon) - rP(r)] \\ & + \alpha' \sum_{j=1}^{N} [K(r+\epsilon, r_j)P(r+\epsilon)P(r_j) - K(r, r_j)P(r)P(r_j)] \end{aligned} \tag{14}$$

where $\alpha' \approx \alpha M$ is the rescaled interaction rate. The mean-field master equation is then obtained by setting, $x = r/\Omega$, and performing a Kramers-Moyal expansion to the lowest order (*5*). For simplicity retaining the symbol $r$ to represent queen gene expression, we find

$$\partial_{\tilde{\tau}} r = \tilde{\alpha}_1(x) = 1 - r - \tilde{\alpha} \sum_{i\neq j} K(r, r_j)P(r_j)\,, \tag{15}$$

where

$$\tilde{\alpha}_1(r) = \Omega^{-1} \int_{-\infty}^{\infty} dr'(r' - r)W(r'|r) \tag{16}$$

is the first jump moment and $\tilde{\alpha} = \alpha'/\Omega^2$. We henceforth refer to the dual version of Eq. (15) describing the evolution of the probability density as the mean field master equation,

$$\partial_{\tilde{T}} P(r) + \partial_r \left[(1-r)P(r)\right] = \tilde{\alpha}\partial_x \left( P(r) \sum_{j=1}^{N} K(r, r_j) P(r_j) \right) . \quad (17)$$

#### 4.1 Continuum limit

We next we take the continuum limit on the number of individuals, $N \to \infty$, to obtain the time evolution of queen gene expression levels in the "tracer" wasp,

$$\partial_t r = 1 - r - \tilde{\alpha} \int_0^\infty K(r, r') f(r') dr' , \quad (18)$$

as well as the time evolution of the population composition, defined by the probability of finding an insects with a given queen gene expression level, $r$,

$$\partial_{\tilde{T}} f(r) + \partial_r \left[(1-r) f(r)\right] = \tilde{\alpha}\partial_r \left( f(r) \int_0^\infty K(r, r') f(r') dr' \right) . \quad (19)$$

This represents the mean-field description of Eq. (9) which is valid in the limit of large populations and time scales, i.e. in the steady state. As we will discuss below, while Eq. (18) and Eq. (19) are not suitable for quantitatively describing the reprogramming dynamics they nevertheless are capable of providing mathematical insight into the mechanisms underlying the regulation of specialisation and plasticity. It is interesting to note that Eq. (19) is conceptually similar to equations described in other biological contexts, such as quorum-sensing bacteria (*6*) or Mitogen competition by stem cells (*7*). In this context, it is also worth to note that in spatially structured systems specialisation can be achieved by spatially separating different phenotypes, such as via the Turing mechanism, spinodal decomposition, lateral inhibition or via external signalling gradients (*8*, *9*). If the spatially homogeneous state is unstable such systems are naturally "plastic".

The dynamics on the molecular scale are coupled through a collision-like functional that represents the effect of repressive interactions. In the next section we will study how such a coupling gives rise to specialization and plasticity as emergent properties of the system.

#### 5 Derivation of the phase portrait

The individual and collective dynamics derived in the previous section provide a mean-field description of the system. In Eq. (19) the time evolution of $f(r, \bar{\tau})$ is governed by a term describing the molecular dynamics (second term on the left hand side) and a term describing the collective behaviour on the population scale (term on the right hand side). The equation gives rise to a steady state when the molecular dynamics is balanced by the population-level feedback. To understand the relaxation dynamics to the steady state, and its stability, it is instructive to consider the co-evolution of the molecular scale, given by the queen gene expression level $r$, and the population scale, represented by the distribution $f(r, t)$. In this section we will illustrate the results of our analysis by means of a phase portrait of the multi-scale dynamics. Although the limits we take, such as taking the mean-field limit, do not accurately reflect the full biological complexity, our approximations are validated by comparison to simulations of the full stochastic dynamics.

The starting point of our analysis are Eq. (18) and Eq. (19), describing the individual and collective dynamics, respectively. Taken together, these equations describe the co-evolution of the queen gene expression level of an individual and the population structure. Stable fixed points of such dynamics represent possible phenotypes in the society, such as queen and workers. We calculate these fixed points from the intersection of the nullclines of the system. These nullclines are given by

$$0 = 1 - r - \tilde{\alpha} \int_{-\infty}^{\infty} K(r, r') f(r') dr' \tag{20}$$

$$\partial_r \left[(1-r) f(r,t)\right] = \tilde{\alpha} \partial_r \left( f(r,t) \int_{-\infty}^{\infty} K(r, r') f(r', t) dr' \right) . \tag{21}$$

Eq. (20) and Eq. (21) provide the basis for understanding the steady state of the system as a function of the population composition and the fixed points of the individual dynamics. Although the population composition is represented by a probability distribution, $f$, its functional nature complicates intuitive interpretations of the relaxation dynamics. In order to obtain a more intuitive picture, we reduce the system to a two-dimensional system describing the coupled evolution of $r$ and the first moment of the population composition, $\langle r \rangle$. The starting point of this approximation is Eq. (18),

$$\partial_{\tilde{\tau}} r = 1 - r - \tilde{\alpha} \int_0^{\infty} r r' \Theta(r' - r) f(r') dr' , \tag{22}$$

describing the evolution of the molecular degree of freedom of a tracer individual. The integral in the right hand side of the equation represents the effect of the interactions received by the tracer individual. In the limit of long times compared with the typical interaction time scale, $t \gg \alpha^{-1}$, the effect of interactions can be approximated by the overall effect of interacting with an effective individual with gene expression level $\langle r \rangle$,

$$\partial_{\tilde{\tau}} r \approx 1 - r - \tilde{\alpha} r \langle r \rangle \Theta(\langle r \rangle - r) , \tag{23}$$

where $\langle r \rangle = \int_0^{\infty} r f(r,t) dr$. Further, by multiplying Eq. (19) by $r$ and integrating over $r$, we obtain the time evolution of the first moment,

$$\partial_{\tilde{\tau}} \langle r \rangle = 1 - \langle r \rangle - \tilde{\alpha} \int_0^{\infty} dr \int_0^{\infty} dr' K(r, r') f(r) f(r') , \tag{24}$$

where $\int_0^\infty dr \int_0^\infty dr' K(r,r')f(r)f(r')$ is the total interaction rate at time $t$ in the nest. To close the system of equations we approximate $\int_0^\infty dr \int_0^\infty dr' K(r,r')f(r)f(r') \approx \langle r\rangle^2/2$ where the factor 2 ensures arises due to double counting of subdominant interactions. Finally, our reduced system of equations reads

$$\partial_{\tilde{T}} r \approx 1 - r - \tilde{\alpha} r \langle r\rangle \Theta(\langle r\rangle - r)\,, \tag{25}$$

$$\partial_{\tilde{T}} \langle r\rangle \approx 1 - \langle r\rangle - \tilde{\alpha}\frac{\langle r\rangle^2}{2}\,. \tag{26}$$

Eq. (25) and Eq. (26) form a two-dimensional system of equations that is amenable for a bidimensional graphical representation given by a phase portrait. The solutions of these equations in the steady state provide the fixed points of the dynamics. In the steady state we find,

$$\langle r\rangle_0 = \frac{\sqrt{2\tilde{\alpha}+1}-1}{\alpha}\,. \tag{27}$$

For $\tilde{\alpha} = 0$ Eq. (25) and Eq. (26) admit only one stable solution corresponding to high queen gene expression levels, $r_0 = 1$ (Figure 2). For $\alpha > 0$, we find three solutions if

$$\langle r\rangle > \frac{\sqrt{4\tilde{\alpha}+1}-1}{2\tilde{\alpha}}\,, \tag{28}$$

which in the steady state is always fulfilled for $\tilde{\alpha} > 0$. These three solution comprise two stable branches, at $r_0 = 1$ and $r_2 = 1/(1 + \tilde{\alpha}\langle r\rangle)$ and an unstable branch at $r = \langle r\rangle$. Formally, the system therefore comprises a saddle node bifurcation with the population structure as a bifurcation parameter. Intrinsic perturbations, which do not change the value of $\langle r\rangle$, are therefore suppressed by the bistable dynamics in the steady state. Substituting the population steady state,$\langle r\rangle_0$ we obtain as intersections

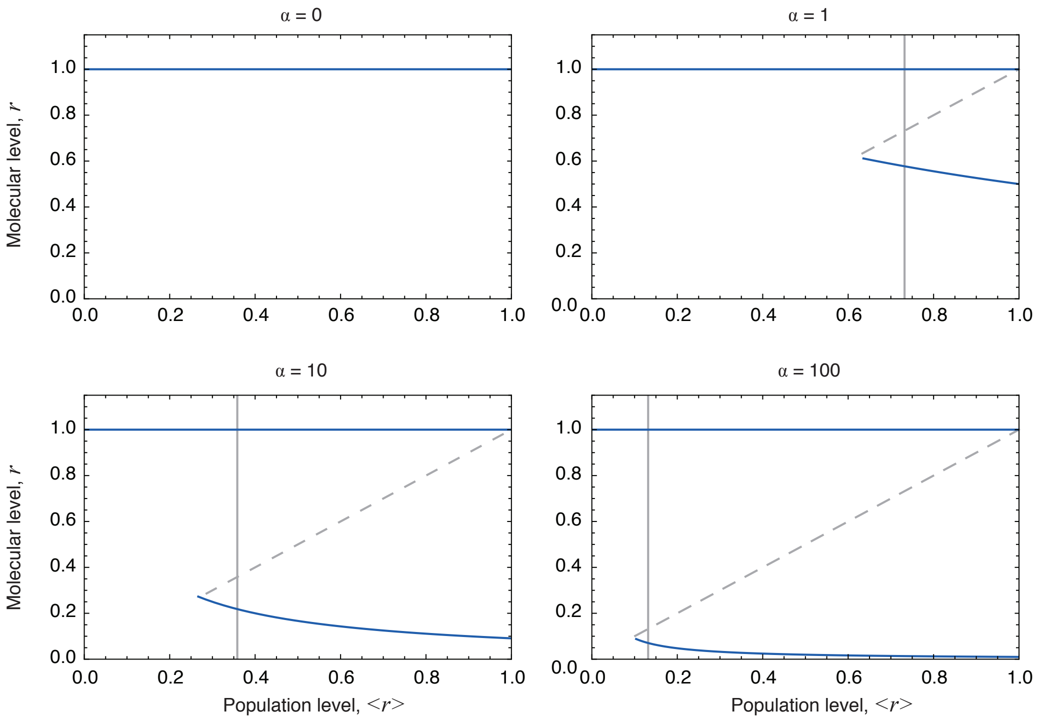


Figure 2: Bifurcation diagram showing fixed points of Eq. (25) as a function of $\langle r\rangle$. Stable branches are represented by solid lines and unstable branches by dashed lines. Vertical lines denote the nullclines of Eq. (26). For $\tilde{\alpha} > 0$ the dynamics undergo a saddle node bifurcation with the population composition, $\langle r\rangle$, as a bifurcation parameter.

of the nullclines for $\tilde{\alpha} > 0$,

$$\begin{aligned} r'_0 &= 1\,, \\ r'_1 &= \frac{\sqrt{2\tilde{\alpha}+1}-1}{\tilde{\alpha}}\,, \\ r'_2 &= \frac{1}{\sqrt{2\tilde{\alpha}+1}}\,. \end{aligned} \tag{29}$$

The flow of the system towards the steady state is represented in Figure 3 and in Figure 3B of the main text. In this representation the $x$ axis corresponds to the molecular degree of freedom and the $y$ axis to the population composition, represented by the first moment of the distribution, $\langle r\rangle$. A population in this portrait is represented

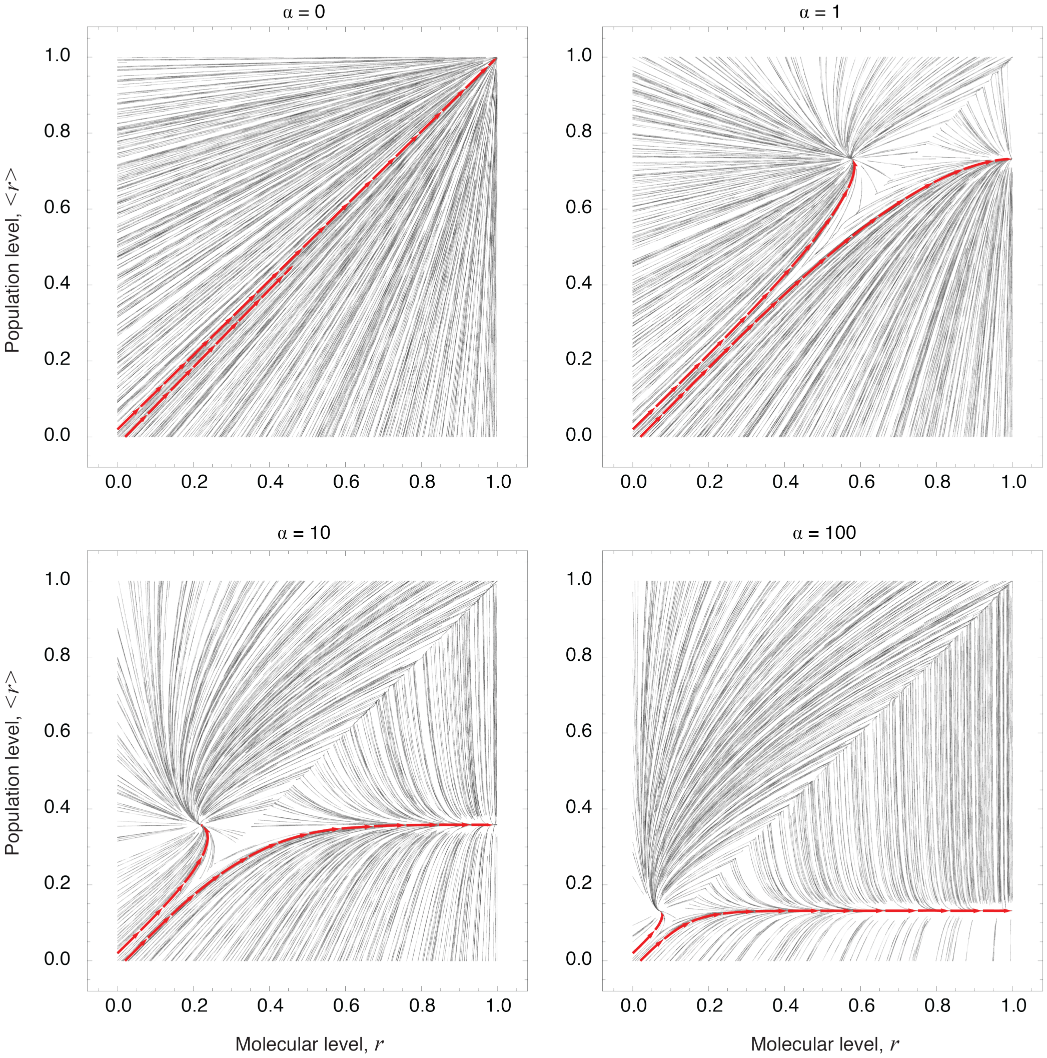


Figure 3: Phase portraits of Eq. (25) and Eq. (26) for different values of $\tilde{\alpha}$. Random trajectories are represented by gray lines and two trajectories originating around the point $(0, 0)$ corresponding to a nest with only individuals lacking queen gene expression are highlighted in red.

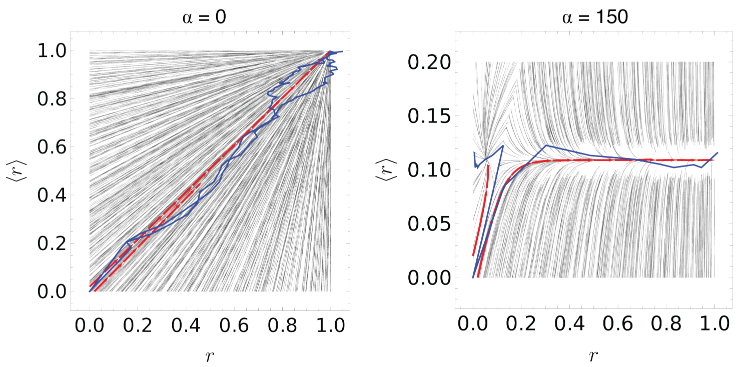


Figure 4: Phase portraits of Eq. (25) and Eq. (26) for different values of $\tilde{\alpha}$. As in Figure 3 random trajectories are represented by gray lines and two trajectories originating around the point $(0, 0)$ corresponding to a nest with only individuals lacking queen gene expression are highlighted in red. Blue lines denote trajectories from stochastic simulation of the full stochastic system, Eq. (9).

by a set of $N$ points, each of them with a different value of the $x$ coordinate, but all of them with the same value of the $y$ coordinate. Changes in the molecular degree of freedom are reflected in changes in the population composition that again affect all individuals identically.

The flow in the mean-field limit qualitatively represents the flow obtained from stochastic simulations of Eq. (9) as shown in Figure 4. Therefore, while our calculations are strictly only valid in the limit of an infinite population size and on long time scales, Eq. (25) and Eq. (26) accurately represent the qualitative structure of the deterministic phase space of Eq. (9).

After the removal of the queen the population obtains a homogeneous structure narrowly distributed around the separatrix defined by $r = \langle r \langle$ and $r = \langle r \langle \ll 1$. From these initial conditions, the dynamics evolve rapidly along the separatrix. On this separatrix fluctuations can drive each individual either to the queen attractor or to the worker attractor.

#### 6 Stability of the steady state

In section 5 we have analyzed the fixed points of the dynamics of the system and studied its stability against uncorrelated and correlated perturbations. To this end, we employed a mean-field approximation effectively neglecting fluctuations arising due to the finite number of individuals and stochasticity in interaction times, Eq. (18) and Eq. (19). We will now address the question of how the stability of the nest and the social order, manifest in a clear separation of queen and worker phenotypes, can be maintained in the presence of strong fluctuations. Indeed, typical population sizes range between 8 to 30 individuals and interactions occur on similar time scales as typical protein degradation times, suggesting that fluctuations are highly relevant for the dynamics. In this section we will ask how the stability of the social structure can be maintained in the presence of strong fluctuations.

To understand how the lifetime of the society is determined by stochastic fluctuations we calculate the probability that an individual upregulates its queen genes in a given time interval. Once a new queen has emerged the society is then destabilized. Therefore, we study the probability of an individual successfully transiting from a worker state to a queen state in a given time interval by producing a sufficient number of queen gene products.

The probability of an individual to increase the number of queen gene products from $r_0$ to the steady state value corresponding to the queen phenotype, $r_Q = 1$, is equal to the probability of such an individual not interacting with the queen during the time that it takes to produce the required number of gene products. Denoting the probability of an individual with $r_i$ gene products not being involved in subdominant interactions during the time interval $\Delta t$ that it takes to produce one more gene product by $\pi_{r_i}$, the probability of an individual successfully upregulating from $r_0$ to

1, $P(1,t|r_0,t_0)$, can be written in the form

$$P(1,t|r_0,t_0) = \prod_k (1-\pi_{r_0+k\epsilon}) = \prod_k [1-\tilde{\alpha}(r_0+k\epsilon)\Delta t] \approx e^{-\sum_k \tilde{\alpha}(r_0+k\epsilon)\Delta t}\,, \tag{30}$$

where we have assumed that the time that it takes to produce a single gene product is deterministically given by its average, $\Delta t$. From Eq. (30) we obtain the probability of observing a successful queen turnover event in a homogeneous population of $N$ individuals with $r_0$ gene products at time $t_0$, in the time $t = (1-r_0)\Delta t$ that it would take to produce $1-r_0$ gene products. This probability is equal to the probability of at least one individual successfully upregulating in this time interval,

$$P(1,t|\{r_0,t_0\}) = 1-(1-P(1,t|r_0,t_0))^N\,, \tag{31}$$

where $P(1,t|r_0,t_0)$ is given by Eq. (30). In order to make the calculation more transparent we explicitly perform the sum over $k$ and approximate $1-e^{-x} \approx x$, obtaining

$$P(1,t|\{r_0,t_0\}) \approx 1-\left(\frac{1}{2}\tilde{\alpha}(1-r_0)(2+r_0)\Delta t\right)^N\,. \tag{32}$$

Replacing $t = (1-r_0)\Delta t$ and given that $r_0 \ll 1$ we finally obtain the probability of observing a queen turnover event in a time interval $t-t_0$,

$$P(1,t|\{r_0,t_0\}) \approx 1-\left(\frac{1}{2}\tilde{\alpha}t\right)^N\,. \tag{33}$$

For typical values of the parameters $N \approx 8-30$ and $\tilde{\alpha}, t \approx 1$, reflecting the fact that molecular and population-level time scales are similar, the probability of observing a queen turnover event is approximately one and, thus, the lifetime of the society is comparable to typical molecular time scales given by the protein degradation time, $\delta^{-1}$. This result, based on a simple comparison of time scales and the constitutive expression of queen genes in the absence of interactions, is in seeming contradiction to the experimentally observed stability of *Polistes* societies over much longer time scales. This raises the question of how the social structure is stabilised. In the remainder of this section we will ask whether the observed reduction of gene expression variance in workers (Fig. 4B of the main text) provides a mechanism for stabilising the society in the long term.

In a given population composed of $N$ workers and one queen the lifetime of the society is dominated by the contribution coming from the working having the highest queen gene expression level. If $r_m$ denotes the maximum queen gene expression level across $N$ workers and if queen gene expression levels in such a population are normally distributed with mean $\mu$ and variance $\sigma^2$, the distribution of $r_m$ is given by the N-th order statistic,

$$P(r_m) = \frac{2^{\frac{1}{2}-N} N e^{-\frac{(r_m-\mu)^2}{2\sigma^2}} \mathrm{Erfc}\left(\frac{\mu-r_m}{\sqrt{2}\sigma}\right)^{N-1}}{\sqrt{\pi}\sigma}\,, \tag{34}$$

where $\mathrm{Erfc}(x) = 1 - \mathrm{Erf}(x)$. We then obtain the probability of a new queen emerging by multiplying the conditional probabilty, $P(1, t|r_m, t_0)$, with $P(r_m)$ and integrating over $r_m$,

$$\int_0^\infty P(1, t|r_m, t_0) P(r_m) dr_m \approx P(1, t|r_m, t_0)|_{r_m^0} \equiv T^{-1}\,, \tag{35}$$

where we performed a saddle-point approximation of the integral around the mode $r_m^0$ of the distribution $P(r_m)$ and $T$ is the lifetime of the society.

For a population distributed normally with mean $\mu$ and variance $\sigma^2$ we find

$$T_{\text{normal}} \approx \exp\left\{-\frac{1}{2}\tilde{\alpha}\Delta t\left[\left(\sqrt{2}\sigma\left((\gamma-1)\mathrm{Erfc}^{-1}\left(\frac{2N}{N+1}\right) - \gamma\mathrm{Erfc}^{-1}\left(2-\frac{2}{eN+e}\right)\right)+\mu\right)^2 - 1\right]\right\}\,, \tag{36}$$

where $\gamma \approx 0.5772$ is the Euler-Mascheroni constant. For an exponentially distributed

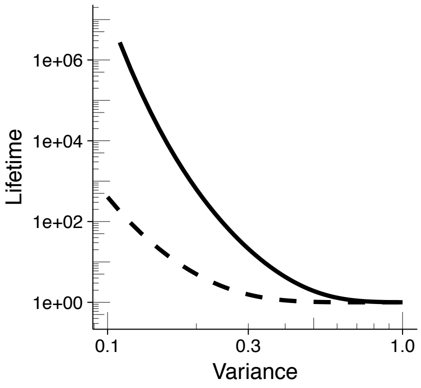


Figure 5: Lifetime of the society as a function of the population variance for normally (solid line) and exponentially (dashed line) distributed populations.

population with mean $1/\sigma^2$ we obtain

$$T_{\mathrm{exp}} = e^{-\frac{1}{2}\tilde{\alpha}\Delta t\left((H_N)\sqrt{\sigma}-1\right)}\,, \tag{37}$$

where $H_N$ is the $N$-th harmonic number.

In order to understand how epigenetic modifications associated with a reduction in gene expression variance alter the stability of the society we investigate how Eq. (35) scales with the variance of queen gene expression across individuals. In Figure 5 the lifetime of the society is shown for exponentially and normally distributed populations. In both cases the lifetime of the society decreases several orders of magnitude as a function of the gene expression variance.

Our theoretical results indicate how epigenetic modifications can increase the lifetime of the society, suppressing noise-induced transitions between different phenotypes. We emphasise that our analysis is independent of the particular molecular mechanism underlying the regulation of gene expression by DNA methylation in *Polistes*. Rather, our results only depend on the empirical observation of reduced gene expression variance between workers for methylated genes (Fig. 4B of he main text).

### 7 Prediction of experimental data

Having derived the model describing the evolution of the joint probability of gene expression and queen gene repressors in Eq. (9), we now set out to fix its parameters and predict the experimental data. To derive a mathematical description that is less dependent on parameters describing poorly understood molecular processes, such as the dynamics of queen gene repressors, we now derive an effective description of the time evolution of the marginal distribution of ovary sizes. To this end, we will derive a description based on ovary growth starting from the model derived above. As ovary sizes are a mere downstream effect of the queen gene expression dynamics (see below), such an effective description should be structurally similar to the master equation describing the time evolution of the marginal distribution of queen gene expression levels, Eq. (13), if the time scales are chosen appropriately.

#### 7.1 Marginal distribution of ovary sizes

In order to predict the time evolution of the probability of observing ovaries of size $\{o_i\}$ at time $t$, $P(\{o_i\}, t)$, we first note that many of the queen genes have functions related to reproduction or are directly responsible for ovary development, such that the instantaneous rate of ovary growth is given by a function that depends on the expression level of queen genes, $g(n_i)$. The function $g(n)$ summarises a cascade of molecular pathways which are not understood in detail. We here make the simplest possible assumption about the functional form of $g(n)$, namely that it depends linearly on gene expression levels and that ovaries grow if the gene expression level exceeds a threshold, $n_0$, and shrink otherwise. Therefore, the rate of ovary growth takes the form $g(n) \propto n - n_0$. As eggs are laid once they have reached a mature size, we impose a reflective boundary condition on $o_i$ at a size $o_0$ which we set to be the maximum ovary size observed in the experiment, $o_0 = 2.5\,\text{mm}$. With this, switching back to dimensional quantities, the time evolution of the conditional probability of ovary sizes follows a master equation of the form

$$\frac{d}{dt}P(\{o_i\},t|\{n_i,q_i\}) = g(n_i)\theta(n_i-n_0)\left[P(\{o_i-1\},t|\{n_i,q_i\}) - P(\{o_i\},t|\{n_i,q_i\})\right] \tag{38}$$
$$- g(n_i)\theta(n_0-n_i)\left[P(\{o_i+1\},t|\{n_i,q_i\}) - P(\{o_i\},t|\{n_i,q_i\})\right] .$$

As, in this model, the expression level of queen genes is independent of ovary size, the joint probability $P(\{n_i,o_i\},t)$ factorizes as $P(\{n_i,o_i\},t) = P(\{o_i\}|\{n_i\},t)P(\{n_i\},t)$. Therefore, the dynamics of the joint probability of $\{n_i,o_i,q_i\}$ are described by

$$\begin{aligned}\frac{d}{dt}P(\{n_i,o_i,q_i\}) = \sum_{i=1}^{N+1}\Big\{&\mu(1-q_i)\left[P(\{n_i-1,o_i,q_i\}) - P(\{n_i,o_i,q_i\})\right] \\ &+ \delta\left[(n_i+1)P(\{n_i+1,o_i,q_i\}) - n_iP(\{n_i,o_i,q_i\})\right] \\ &+ g(n_i)\theta(n_i-n_0)\left[P(\{n_i,o_i-1,q_i\}) - P(\{n_i,o_i,q_i\})\right] \\ &- g(n_i)\theta(n_0-n_i)\left[P(\{n_i,o_i+1,q_i\}) - P(\{n_i,o_i,q_i\})\right] \\ &+ \Gamma(t_i^{\text{int}})P(\{n_i,o_i,1\})(1-2q_i) \\ &+ \omega\sum_{j\neq i}K(n_i,n_j)P(\{n_i,o_i,0\})P(\{n_j,o_j,q_j\})(2q_i-1)\Big\} ,\end{aligned} \tag{39}$$

with $g(n)$ denoting the rate of ovary growth for a gene expression level $n$.

With the aim of comparing our simulation results to the experimental ovary dissection data we first integrate out gene expression, yielding an equation describing the evolution of ovary sizes. If the persistence time of the repressive effect of interactions plus the typical production time of gene products is smaller than the typical time between two interactions, individuals alternate periods of growing and shrinking of their ovaries with the duration of these periods determined by the ratio between the typical interaction and persistence times plus the queen gene production times. Taking this limit, the ovary growth rate only depends on whether queen gene expression is above or below the threshold $n_0$. For that purpose we define a random variable,

$$s_i = \Theta(n_i - n_0)\,, \tag{40}$$

such that the ovary growth rate is proportional to $2s_i - 1$.

The time evolution of the random variable $s_i$ is linked to the dynamics of $n_i$. This gives rise to an explicit time delay in the ovary equation. Specifically, following an interaction, an individual with $s_i = 1$ needs a time $t_{\rm off}$ to degrade enough gene products and activate the pathways responsible for flipping the ovary growth state, $s_i = 0$. On the other hand, if an individual with $s_i = 0$ does not engage in a subdominant interaction during a time $t_{\rm on} \approx t_{\rm off} + t_{\rm per}$, given by the sum of the persistence time of queen gene repressors and the time needed to express queen genes beyond a level $n_0$ and activate pathways related to reproduction, it will again flip the ovary growth state, $s_i = 1$.

In the long time limit the ovary size of a given individual is determined by the sign of the diference $T_{s_i=1} - T_{s_i=0}$ where $T_{s_i=j} = \int_0^t dt'\theta(s_i - j)$ is the time that the ovary growth rate spends in the state $j$. In this limit, the instantaneous value of gene expression $r$ at time $t$ is a good indicator of the sign of $T_{s_i=1} - T_{s_i=0}$, and hence of ovary growth. Taking this into account the interaction kernel can be expressed in terms of ovary size and we obtain the effective master equation governing the evolution of the marginal distribution of ovary sizes,

$$\frac{d}{dt}P(\{o_i, s_i\}) = \sum_{i=1}^{N+1}\Big\{g(s_i)\left[P(\{o_i - 1, s_i\}) - P(\{o_i, s_i\})\right] + \delta(t_i^{\text{int}} - t_{\text{off}})P(\{o_i, 1\})(1 - 2s_i) + \delta(t_i^{\text{int}} - t_{\text{on}})P(\{o_i, 0\})(2s_i - 1)\Big\}, \tag{41}$$

where $t_i^{\text{int}}$ is, as before, the time elapsed since the last subdominant interaction of individual $i$. Eq. (41) provides a description of the system at the ovary level that retains the foremost characteristics present in the experimental data, i.e. the presence of multiple queens in the nest shortly after reprogramming and the posterior relaxation towards an analogous state to the control, as demonstrated by the results in Figure 2F of the main text. The existence of explicit time delays $t_{\text{on}}$ and $t_{\text{off}}$ is responsible for the transient observation of multiple queens during reprogramming. Specifically, such an overshoot arises if the value of $t_{\text{off}}$ is of similar magnitude as the time scale associated with ovary growth.

#### 7.2 Experimental parameters

The model defined in Eq. (41) includes several parameters. In this section we provide justifications for the parameter values used to predict the experimental observables in Fig. 2F and Fig. S2F of the main text. In the derivation of the model, Eq. (41), we did not assume any non-linearities unless supported by experimental data. Such non-linearities, for example in the relation between ovary growth and gene expression, naturally exist in any biological system. Therefore, we have to exercise some caution in interpreting these parameters in literal biological terms. We still expect, however, that the order of magnitude of parameters is not altered by unknown non-linearities

| Parameter | Value | Justification |
|---|---|---|
| $\alpha$ (Interaction rate) | 1 $\text{day}^{-1}$ | Measurement from video recordings presented in Fig. 2B of the main text. |
| $\tau_{\text{off}}$ (Typical time between an interaction and changes in ovary growth) | 2.5 days | We estimated this parameter based on the observation of 2-3 egg layers in the early commitment phase. |
| $o_r$ (Ovary growth rate) | $0.25\,\text{mm}\,\text{day}^{-1}$ | The first egg layer is observed 6 days after queen removal with a size of mature eggs of 1.5 mm. |

Table 1: Summary of parameter values used for predicting experimental data.

and can be estimated by independent observations from experiments or the literature. Parameter values are summarised in Table 1.

The time evolution of the global activity in a nest (Fig. S2F) is determined by a component which is independent of the fighting interactions involved in the regulation of the reprogramming process and a component reflecting these interactions. Only the latter component is reflected in the model. We note that this component is proportional to the total interaction rate. From Eq. (41) we find that this interaction rate is

$$\sum_{ij} o_i o_j f(o_i) f(o_j) = \left( \sum_i o_i f(o_i) \right) \left( \sum_j o_j f(o_j) \right) = \langle o_i \rangle^2 .$$

In Fig. S2F, to take into account the different contributions to the global activity mentioned above, an we added an offset value to the theoretical prediction such that the empirical and theoretical values matched in the control phase. Then, activity levels were rescaled such that the maximum of both curves matched.

#### 8 Numerical simulations

We performed kinetic Monte Carlo simulations following Gillespie's algorithm to obtain approximate solutions to master equation for a population of $N$ individuals (*10*). Out of the different possible processes - production of a new gene product, degradation of gene products, interactions, degradation of the queen gene repressors or ovary development - one was randomly selected with probability proportional to the overall rate of the respective process (in vector form)

$$\left(\mu, \delta n_i, \alpha K(n_i, n_j), \delta(t^i_{int} - t_{per}), g(n_i)\right) \ . \tag{42}$$

Once a process has been selected the state of the system is updated depending on the selected reaction and finally, the simulation time is advanced an amount $\Delta t$ drawn from an exponential distribution of parameter $\lambda$ given by the inverse sum of the overall rates,

$$\lambda = \left\{ \sum_i \left( \mu + \delta n_i + \sum_j K(n_i, n_j) + \delta(t^i_{int} - t_{per}) + g(n_i) \right) \right\}^{-1} \ . \tag{43}$$

Unless specified otherwise the following parameters were used for all the simulations $\mu = 500$, $\delta = 1$, $n_0 = 250$, $\lambda = 10$, $t_{per} = 1$. The ovary growth rate was chosen so that ovaries were mature in 6 days, as observed experimentally.

All simulations were implemented in Julia and the source code is available upon request to the authors.
